# Identifying signaling genes in spatial single cell expression data

**DOI:** 10.1101/2020.07.27.221465

**Authors:** Dongshunyi Li, Jun Ding, Ziv Bar-Joseph

## Abstract

**Motivation:** Recent technological advances enable the profiling of spatial single cell expression data. Such data presents a unique opportunity to study cell-cell interactions and the signaling genes that mediate them. However, most current methods for the analysis of this data focus on unsupervised descriptive modeling, making it hard to identify key signaling genes and quantitatively assess their impact.

**Results:** We developed a **M**ixture of **E**xperts for **S**patial **S**ignaling genes **I**dentification (MESSI) method to identify active signaling genes within and between cells. The mixture of experts strategy enables MESSI to subdivide cells into subtypes. MESSI relies on multi-task learning using information from neighboring cells to improve the prediction of response genes within a cell. Applying the methods to three spatial single cell expression datasets, we show that MESSI accurately predicts the levels of response genes, improving upon prior methods and provides useful biological insights about key signaling genes and subtypes of excitatory neuron cells.

**Availability:** MESSI is available at: https://github.com/doraadong/MESSI

## 1 Introduction

The ability to determine gene expression at the single cell level led to several novel findings and insights (Tanay and Regev, 2017; Cembrowski, 2019). Recently, a number of new technologies take this a step further: enabling the profiling of spatial single cell expression (Moffitt *et al*., 2018; Wang *et al*., 2018). These techniques, which primarily rely on combinatorial in situ hybridization (FISH) or in situ sequencing, can determine the levels of hundreds to thousands of genes for each cell while still retaining information of the location of the cells. Such information opens the door to much more detailed and accurate analysis of the impact of spatial location on cell-cell interactions and gene expression (Lein *et al*., 2017).

There have been a number of methods that attempted to infer cell-cell interactions from scRNA-Seq data. Since such data does not contain spatial information, these methods have mainly focused on the co-expressed ligand-receptor pairs between cell types (Cabello-Aguilar *et al*., 2020; Efremova *et al*., 2020; Wang *et al*., 2019; Tsuyuzaki *et al*., 2019; Kumar *et al*., 2018; Zhou *et al*., 2017). However, given the data used, it was hard to determine if the cell types identified as interacting were indeed close in 2D or 3D space. A number of computational methods have been proposed to analyze and model spatial single cell data. To date, most methods for the analysis of such data are unsupervised focusing on clustering or representation to infer spatial patterns for cells, genes and transcripts (Zhu *et al*., 2018; Edsgärd *et al*., 2018; Svensson *et al*., 2018; Sun *et al*., 2020). Very few methods intended to study interactions between cells using spatial single cell expression data. For example, Giotto (Dries *et al*., 2019) identifies potential interacting neighboring cell type pairs by testing if genes in a particular cell type are differentially expressed when adjacent to another cell type. While Giotto provides some information on the set of interacting genes and proteins, these models are descriptive rather than predictive. Due to the lack of ground truth regarding ligand-receptor interactions in specific tissues or cell types, it is hard to evaluate the accuracy of such methods in a comprehensive manner and to determine whether they indeed capture the true interactions or instead just overfit the observed expression values.

To better infer the set of signaling genes within and between neighboring cells, we developed a framework, **M**ixture of **E**xperts for **S**patial **S**ignaling genes **I**dentification (MESSI), that utilizes Mixture of Experts (MoE) and multi-task learning to jointly model the interactions within and between cells from spatial single cell data. For each cell type in the data, the MESSI model uses as input a subset of inter- and intra-signaling genes to predict the expression of a set of response genes. As we show, a major advantage of the MoE framework for modeling single cell data is its ability to account for subtypes. The use of multi-task learning further enables the sharing of information among response genes via joint learning of response genes’ covariance matrices.

To test our models, we performed cross-validation analysis in which we learned on a subset of the data (in our case, a subset of the animals profiled in different studies) and tested on other animals. Once confirmed, the coefficients assigned by the model to the different signaling genes are analyzed to identify key signaling genes that mediate cell-cell communication and to identify cell subtypes that differ in these and their impact on the response genes’ expression.

We applied our MESSI framework to three spatial single cell datasets. As we show, the method can accurately predict gene expression by utilizing neighboring cells’ information and improves upon several other methods suggested for expression prediction in spatial and non-spatial analysis settings. Analysis of the resulting experts for one of the datasets identifies an important signaling gene that cannot be determined based on expression alone, and that can be used to define an important excitatory neuron subtype.

## 2 Materials and Methods

### 2.1 Datasets and data pre-processing

We tested MESSI using three spatial transcriptomics datasets: the MERFISH hypothalamus data, the MERFISH U-2 OS data, and the STARmap mPFC data. The MERFISH hypothalamus data (Moffitt *et al*., 2018) profiled ~11 million cells in the preoptic region of the mouse hypothalamus. For each cell, the expression for 155 genes was measured using a single-molecule imaging method based on combinatorial or sequential FISH labeling. Multiple animals under different behavioral conditions were profiled, each with roughly 20K to 70K cells. In our analysis, we used all genes profiled and all naive (4), parenting (4), and virgin parenting (5) female animals that were studied.

The MERFISH U-2 OS data (Xia *et al*., 2019) is a single cell-line dataset profiling 1368 cells in three batches. For each cell, the expression of 10050 genes was measured. We selected the most significantly DE (differentially expressed) genes in the clusters identified by Xia *et al*. (2019) for the MESSI modeling resulting in 742 DE genes. The STARmap data includes cells from the mouse medial prefrontal cortex (mPFC) profiled using an in situ sequencing method (Wang *et al*., 2018). We used the three control samples profiled by STARmap, each containing between 1100 and 1300 cells. STARmap measured the values of 166 genes and we used all of them. See *Supplementary Methods* for details on how the data was processed and genes were selected.

To identify potential signaling genes, we used genes determined to be ligands or receptors, including those from Ramilowski *et al*. (2015). See *Supplementary Methods* and Table S1 for detailed information on the set of genes we used. To identify neighboring cells, we used Delaunay triangulation with a specific distance cutoff (for example, 100 micrometers for the MERFISH hypothalamus dataset, Figure 1).

**Fig. 1.**
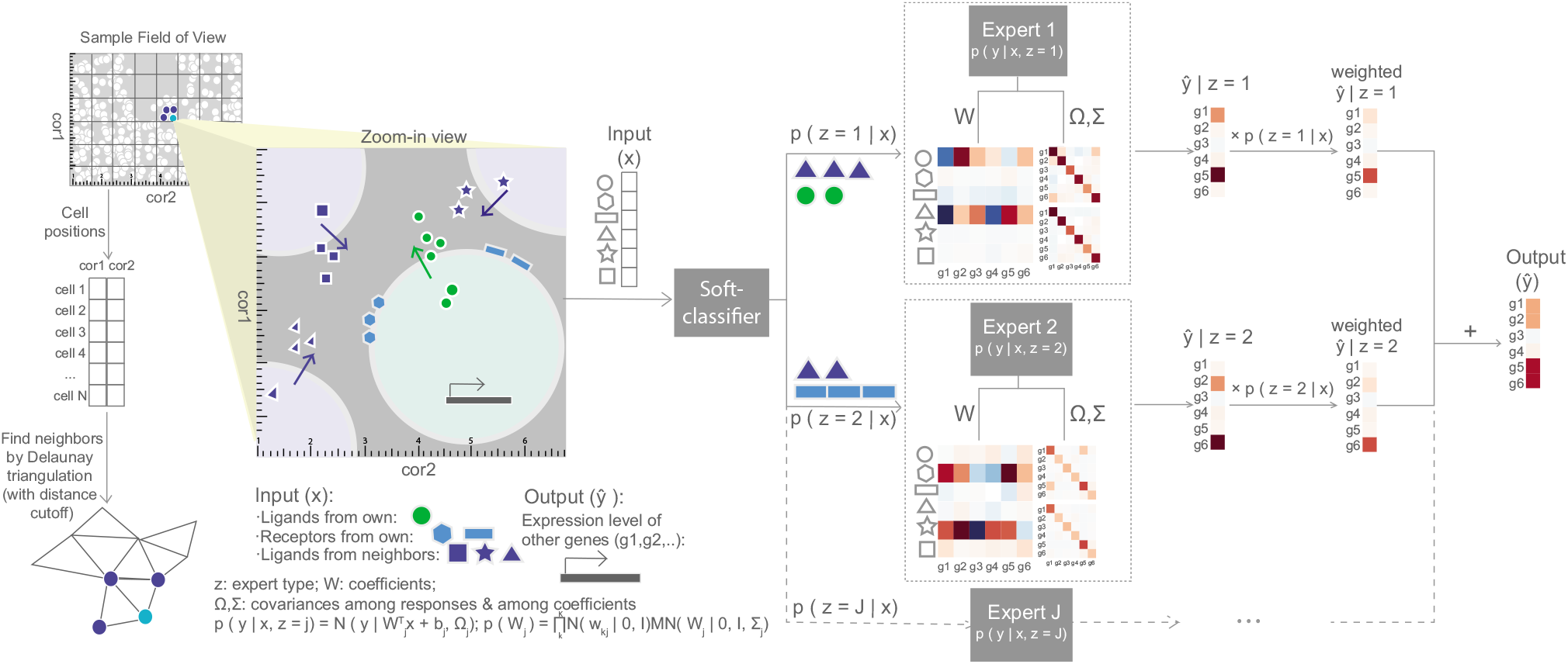
MESSI framework. Cell neighbors are determined by applying Delaunay triangulation to the cell spatial coordinates. Next, expression levels of signaling genes in the cell and its neighbors, the cell’s spatial location, and the neighbors’ cell types, are used to probabilistically assign the cell to a subtype (expert). Experts then integrate information from intra- and inter-signaling genes to predict the expression of response genes. The final output is then the average of these predictions weighted by the expert likelihood.

### 2.2 Inferring inter- and intra-gene-gene interactions

Our goal is to identify genes encoding signaling molecules (termed signaling genes) used for cell-cell interactions. For this, we developed a Mixture of Experts for Spatial Signaling genes Identification (MESSI) framework that aims to predict the expression of genes (termed response genes) in a specific cell type using a combination of intra-cellular signaling molecules, specifically the ligands or receptors produced by the cell itself, and the expression of ligands in neighboring cells (also referred as the intercellular signaling genes). Figure 1 provides a high-level overview of our MESSI framework. Our method utilizes the expression of genes in neighboring cells (dark blue in the zoom-in view of Figure 1) and the expression of genes within a cell (green and light blue) to predict a subset of response genes. Predictions are performed by a set of experts that correspond to potential subtypes of cells learned by the model. Cell assignment to subtypes (experts) is determined by the signaling genes and the additional spatial information (e.g., neighboring cell types and spatial locations). Below we discuss in detail how to formulate the MESSI optimization problem, how to learn parameters for the model, and how we evaluated its performance and extracted biological insights from the model learned.

### 2.3 Mixture of Experts for Spatial Signaling genes Identification(MESSI) framework

We employed and extended a Mixture of Experts (MoE) framework proposed by Jacob and Jordan (Jordan and Jacobs, 1994) for spatial expression prediction. Denote the response gene expression vector by **y** (with *K* entries where *K* is the number of response genes to be predicted). Denote the input features as **x**, which is a *D* dimensional vector (so *D* is the total number of input features). For each cell *i*, we denote by **y_i_** the expression values of its response genes and by **x_i_** the features for this cell. The features we consider are the expression of ligands and receptors in cell i, the expression of ligands in cell *i*’s neighbors, cell *i*’s spatial coordinates (location within the image/tissue) and cell types of its neighbors (we used the cell type annotations from the original studies). Given these definitions, for *N* cells, we construct the output *Y*(*N* × *K*) and input *X*(*N* × *D*) matrices. Assuming that given the expression of ligands in neighboring cells, and the expression of signaling genes within the cell, the expression of response genes in a cell is independent of other cells, we can write the total conditional likelihood as 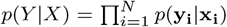. MoE models assume that these conditional distributions are mixtures of conditional distributions produced by different experts (in our case, cell “subtypes”). For each cell *i*, we use *z_i_* as a categorical variable to indicate the expert (“subtype”) that generates cell *i*. Under the mixture assumption of MoE, we have for each cell *i*

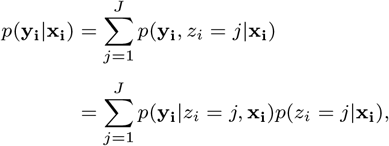

where *J* is the number of experts and *j* is the *j*th expert. In other words, we can express the conditional likelihood as a weighted average of conditional likelihoods *p*(**y_i_**|*z_i_* = *j*, **x_i_**) produced by each *j*th expert. Note that here the contribution (weight) from expert *j* for cell *i*, *p*(*z_i_* = *j*|**x_i_**), is also dependent on the feature variables **x_i_**. This contribution is determined by a classifier (gate).

#### 2.3.1 Multi-output learning model for experts

Given that we are trying to predict multiple genes, we use multi-output models for each expert. These models take into account the dependency among different tasks and so are more suitable for predicting the expression of multiple genes, some of which are likely co-expressed or co-regulated, than methods assuming conditional independence. For this, we extended the MoE framework such that each expert uses a weighted version of the Multiple-output Regression with Output and Task Structures (MROTS) model (Rai *et al*., 2012). For a single cell *i*, the conditional likelihood described by an expert *j*, as modeled by MROTS, is

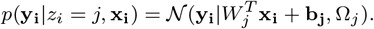

In other words, it assumes a multivariate normal distribution with 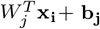 as the mean and Ω*_j_* as the covariance. Here *W_j_*(*D* × *K*) and **b_j_** (*K* × 1) are the model coefficients and intercepts. In addition, the model also assumes a prior on the distribution of the coefficients *W_j_*

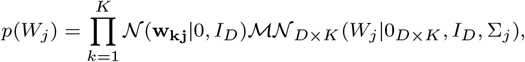

where 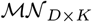 denotes the matrix normal distribution. This distribution for *W_j_* couples the coefficients of different tasks by assuming a column-covariance matrix Σ*_j_*(*K* × *K*) among tasks. The other component of the prior, 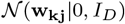, can be viewed as the *l*_2_ regularization on the weights **w_kj_** for each individual task *k*. The covariance matrices Ω*_j_* and Σ*_j_* each characterizes different aspects of the dependence among tasks, Ω*_j_* for the conditional covariance among the outputs and Σ*_j_* for the marginal covariance among the coefficients.

### 2.4 Learning and inference for MESSI

Learning for MESSI is performed using an Expectation–maximization (EM) algorithm which is commonly applied to Mixture of Experts (MoE) models (Jordan and Jacobs, 1994). The algorithm aims to maximize

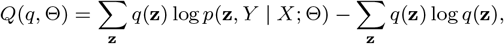

a lower bound of the log conditional likelihood log *p*(*Y*|*X*). Here, **z** is the membership vector of size *N* × 1, *q* is the unknown distribution of **z** and Θ is the collection of all model parameters. Since the distribution *q* of **z** is unknown, in the E-step of iteration *t*, for each cell *i*, we first postulate a distribution for *z_i_* based on the parameters Θ^*t*–1^ learned from the previous iteration. See *Supplementary Methods* for the complete derivation.

Next, we perform the M-step where we maximize *Q*(*q^t^*, Θ) w.r.t. Θ. Given the modularity of the MoE framework, the M-step can be subdivided into separate learning tasks for each expert and classifier (See *Supplementary Methods*). For each expert, we derived and implemented an alternating minimization algorithm for a weighted version of MROTS with *h_j_*(*i*) as the weight of expert *j* for cell *i*. We use logistic regression for the classifier. The learning algorithm iterates between the E-step and M-step until convergence. For a new sample 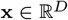, the prediction 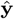 is based on the weighted average of the predictions made by each single experts. The weight is given by the trained classifier based on the input **x**, indicating how likely that the new sample comes from an expert. See *Supplementary Methods* for the detailed derivation of the training algorithm and how the model makes predictions.

To learn hyper-parameters, for example, the number of experts, we employed a nested cross-validation (CV) strategy where the inner loop selects the best model, and the outer loop evaluates the performance of the model on a left-out animal/replicate. For each iteration of the inner loop, we split the training data into a training set and a validation set and conduct a grid search for the values of the hyperparameters using the validation set. After looping through all the training and validation sets pairs, we select the best values for the hyperparameters using majority vote. We then retrain using the entire training dataset with the selected combination of hyperparameter values, and the trained model is then applied to the left-out data. See *Supplementary Methods* for more details.

### 2.5 Baseline and comparison methods used to benchmark MESSI

The baseline model we compared to only uses information about the location of the cell and its type to predict response gene expression. Thus, it does not use any information from neighbor cells. All other methods we compared to use the same set of input features as MESSI. Chen *et al*. (2016) described that researchers in LINCS program (http://www.lincsproject.org/) predicted gene expressions by building a linear regression model for each response gene assuming response genes are conditionally independent. XGBoost (Li *et al*., 2019; Chen and Guestrin, 2016) uses a boosting idea for the prediction. Chen *et al*. (2016) learned multi-layer perceptron (MLP) models. We consider both single- and multi-node output MLP models, where the latter enables learning across response genes. See *Supplementary Methods* for the implementation details of these methods.

## 3 Results

### 3.1 MESSI achieved high prediction accuracy compared to the baseline and comparison methods

We first tested our model on the MERFISH data of the mouse hypothalamus preoptic region (Moffitt *et al*., 2018), where for each cell, the number of RNA molecules of a set of marker genes was measured by multiplexed error robust fluorescence in situ hybridization (MERFISH). The MERFISH hypothalamus data contains ~11 million cells, each with 155 genes profiled from multiple animals in both sexes under naive and several different behavioral conditions, see *Materials and Methods* for details. To determine the ability of our method to accurately predict genes’ expression, we performed nested cross-validation (CV) analysis to learn model parameters and hyper-parameters (see Table S3 for the selected hyperparameter values). We next used the learned model to predict each cell’s expression in the samples of the left-out animal and compared the predicted results to the actual expression levels to determine the accuracy of the model. We compared MESSI’s prediction accuracy with a baseline model that only uses cell types and spatial locations of the cells, but not the expression of genes. We also compared MESSI to regression models that have been previously used to predict gene expression based on a subset of genes for non-spatial data. These include linear regression (LR), multilayer perceptron (MLP) with single- or multi-node output (Chen *et al*., 2016) and XGBoost which uses a boosting algorithm to learn regression trees (Li *et al*., 2019; Chen and Guestrin, 2016). Finally, we compared MESSI to the Multiple-output Regression with Output and Task Structures (MROTS) model (Rai *et al*., 2012), which, unlike MESSI, uses a single expert rather than a Mixture of Experts.

Results are presented in Figure 2. As can be seen, MESSI performed best, achieving the lowest mean absolute error (MAE) when averaged over cells from eight cell types (median of MAE: MESSI: 0.3568 vs. 0.3596 for XGBoost and 0.3607 for MLP single, the next 3 performed worse). Cell type specific results are shown in Figure 3. Again, MESSI performed best for the four major cell types. For example, for excitatory neurons, the median of MAE for MESSI is 0.3757 vs. 0.3824 for XGBoost and 0.3801 for MLP single. These improvements over other models are significant, as shown in Figure S12 and S13. In contrast, the baseline model performed the worst for all cell types (the median of MAE averaged over eight cell types is 0.5477). This indicates that while cell type and location are important features, the specific expression of genes in a cell is significantly affected by intra- and inter-signaling molecules. See Figures S18-S21 for the other cell types.

**Fig. 2.**
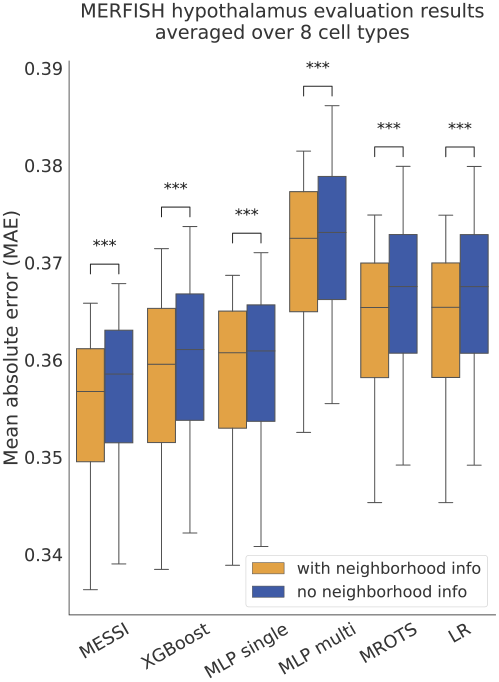
Cross-validation results averaged over cell types. MESSI achieved the lowest mean absolute error (MAE) when averaged over eight cell types from the MERFISH hypothalamus data. For all methods, models utilizing neighborhood expression values significantly outperformed those that do not. with neighborhood info: using both intra- and inter-cellular signaling genes, and other spatial information (see Materials and Methods) as features; no neighborhood info: using only intra-cellular signaling genes as features. ***: p-value below 1e-3; LR: Linear regression; MLP single: MLP with single-node output; MLP multi: MLP with multi-node output; MROTS: Multiple-output Regression with Output and Task Structures.

**Fig. 3.**
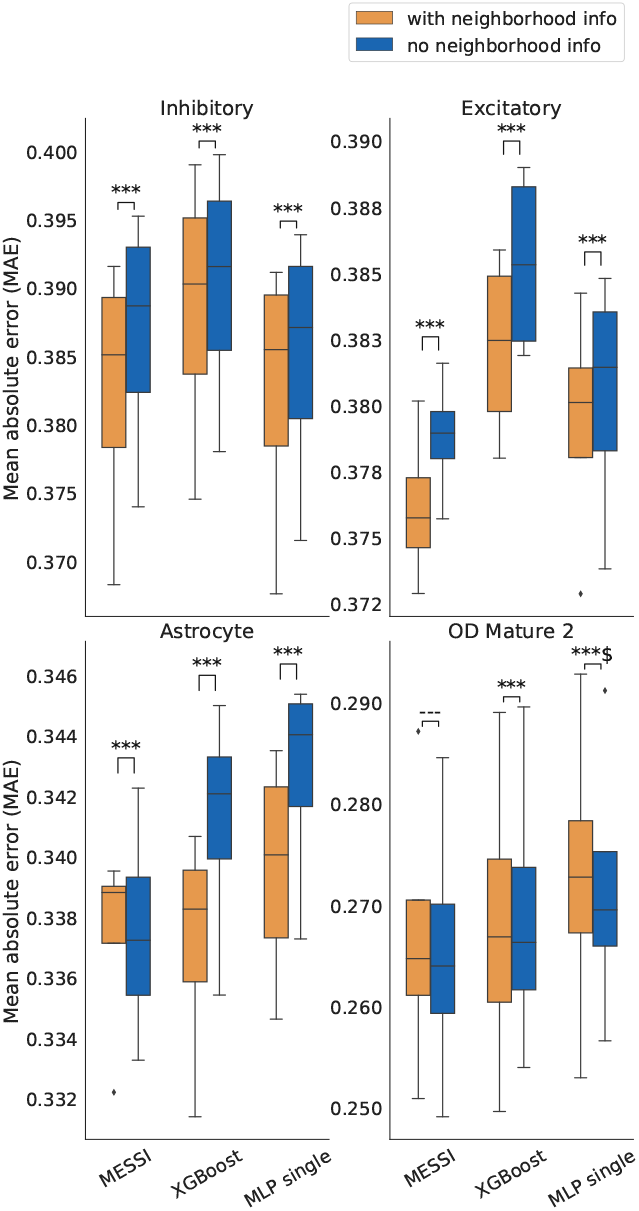
Advantage of using neighborhood expression values. Using both intra- and inter-(neighborhood) cellular features improved prediction accuracy across most major cell types. Here the cell types are listed in the order of decreasing sample size. with neighborhood info: using both intra- and inter-cellular signaling genes, and other spatial information (see Materials and Methods) as features; no neighborhood info: using only intra-cellular signaling genes as features. ***: p-value below 1e-3; ---: p-value larger than 5e-2; $: more than half of the CV groups show non-significant improvement when using neighborhood information; MLP single: MLP with single-node output

To test the impact of neighboring cells, we repeated the analysis above using the same methods, but this time only relying on the expression of intra-cellular signaling genes (i.e., genes in the same cell). Comparison between a model that uses neighboring cell expression levels and one that does not use them is presented in Figures 2 and Figure 3. As can be seen, the application of models that ignore the expression of signaling genes in neighboring cells resulted in significantly higher MAE for all methods. Similarly, all top-performing methods showed improved prediction performance when using inter-cellular signaling genes for most of the major cell types (Figure 3). For example, for excitatory neurons, the median of MAE using neighborhood genes versus not is 0.3757 versus 0.3789 for MESSI, 0.3824 versus 0.3853 for XGBoost, and 0.3801 versus 0.3814 for MLP single.

### 3.2 Impact of sample size on performance

While MESSI performed well on cell types for which many cells were profiled (for example, for the 55K inhibitory cells), it was not the top performer for other cell types (for example, the 6K endothelial 1 cells). We hypothesized that the reduction in performance was not due to the cell type identity but rather to the lack of data, which may lead to overfitting due to the increase in the number of parameters being fitted by MESSI. To test this, we down-sampled the largest two cell types profiled (inhibitory and excitatory) and compared the performance when training on a reduced number of cells to the performance discussed above when using the full set for training. In the smaller training sets, we used a training sample size ranging from 0.7K to 6K compared to 19K to 55K for the full set. Results for such down-sampling are presented in Figure 4. While MESSI outperformed XGBoost when using all available data (Figure 3), when using the reduced dataset, XGBoost was better. More generally, using more samples resulted in an obvious decrease in prediction error for all methods, except for the baseline model. Thus, when enough cells are sampled, models that can take into account the heterogeneity within cell types perform better than simple regression models relying only on cell types and spatial locations.

**Fig. 4.**
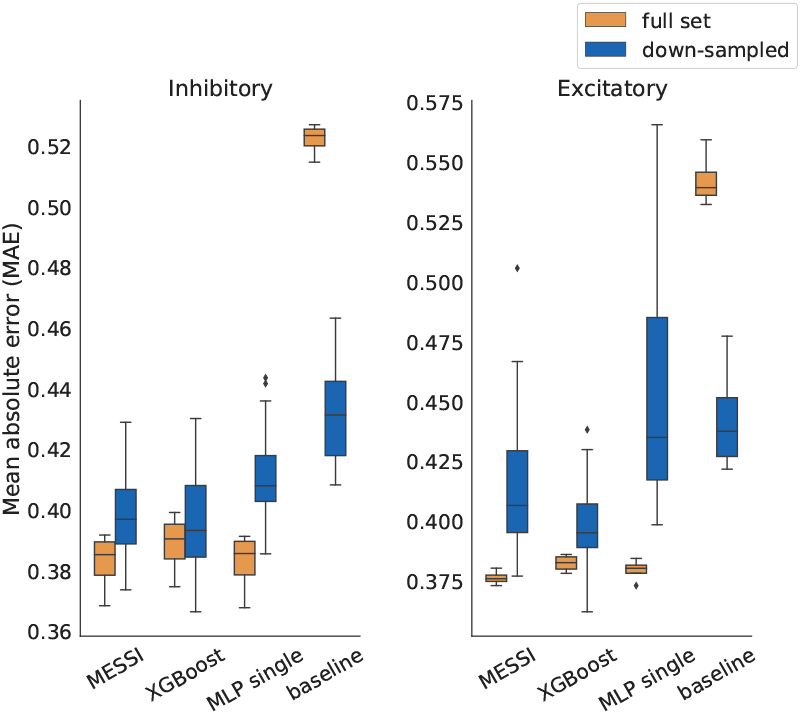
Impact of training sample size on accuracy. We down-sampled inhibitory and excitatory cells to compare the performance of methods when using the full data and when using a smaller dataset. While MESSI and MLP single performed better than XGBoost when using all available data, using the reduced dataset for training likely led to overfitting for these two methods. MLP single: MLP with single-node output

### 3.3 Testing MESSI on additional datasets

To test the generality of our method, we further tested MESSI on two other spatial single cell expression datasets, both with a smaller sample size. The first is another dataset profiled by MERFISH that only focuses on a single cell type (U-2 OS cell line) (Xia *et al*., 2019). This dataset includes about 1.3K cells though many more genes (about 10K) were profiled for each cell. The second is from a method called STARmap, which profiled mouse medial prefrontal cortex (mPEC) cells (Wang *et al*., 2018). Again, this dataset is much smaller than the first MERFISH dataset, with about 2K cells profiled from the biggest cell type, excitatory neurons. STARmap provides information on the expression of 166 genes per cell. Since we use the prediction accuracy of response genes to validate the signaling genes and subtypes MESSI learned, we have also assessed MESSI’s performance when using different sizes of response genes sets. For this, we used the smaller MERFISH data and varied the number of top DE genes selected as response genes between 38 and 693. The nested CV resulted in the selection of a single expert (essentially equivalent to the MROTS method) for all responses sets. Same as for the first MERFISH data, MESSI performed best for this data as well, as shown in Figure 5 when using 38 responses and Figure S7 for all settings (see *Supplementary Results* 2.1 for details). MLP single-node output with the number of hidden nodes equal to 2 performed much worse than the other models and are not shown in these plots (see *Supplementary Results* Figure S10). For the STARmap data, which is also very small, we observed that MESSI again selected only one expert. MESSI’s performance on this data was slightly worse than XGBoost likely due to overfitting given the small training set size (Figure 5). Another potential reason for overfitting the STARmap dataset is the high drop-out rate (on average, 54% of cells having 0 values for a single gene compared to only 13% for the MERFISH U-2 OS cell line data).

**Fig. 5.**
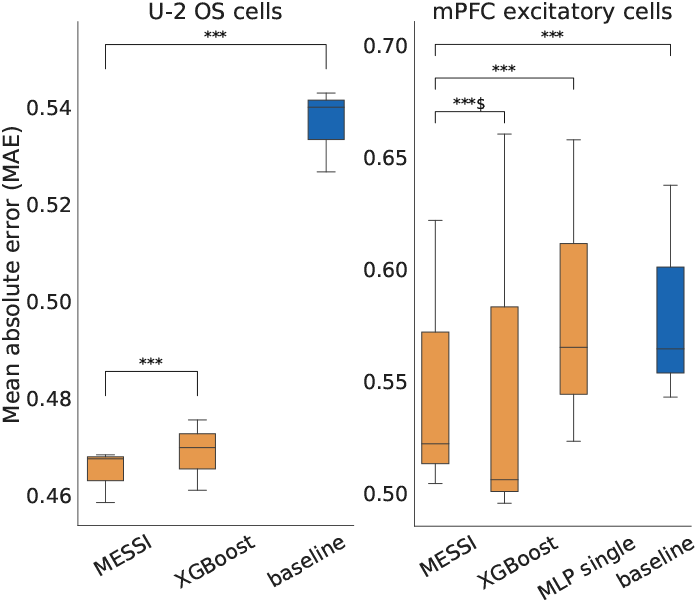
CV results of the MERFISH U-2 OS cell line and the STARmap data. Left: Results for the analysis of the MERFISH U-2 OS cells data. Right: Results for the analysis of the STARmap data. ***: p-value below 1e-3; $: more than half of the CV groups show nonsignificant improvement when using MESSI; MLP single: MLP with single-node output

### 3.4 MESSI identified functional changes in neuron subtypes between conditions

Given MESSI’s ability to accurately predict gene expression based on signaling genes, we used MESSI to explore cell type specific functional changes in signaling networks related to behavioral changes. For this, we compared MESSI models learned from naive female animals, with models learned from virgin females and mothers subjected to behavior stimuli as described in Moffitt *et al*. (2018). Following Moffitt *et al*. (2018), we refer to these samples as virgin parenting and parenting samples.

We first compared the performance of models learned using the naive or behavioral training samples on the task of predicting expression values for unseen behavioral samples. For cell types where the performance of these models is roughly the same, we can conclude that no major changes occur between the two conditions (since models trained on one perform well on the other). However, if we observe large differences in prediction abilities, then for these cell types, signaling molecules’ impact on the expression levels of the response genes are likely different. When using this approach for the MERFISH hypothalamus data, we observed a large improvement in the prediction ability for models learned using behavioral samples for inhibitory and excitatory cell types, while no obvious change in performance for the other cell types (Figure 6 and *Supplementary Figures* Figure S1 & S2).

**Fig. 6.**
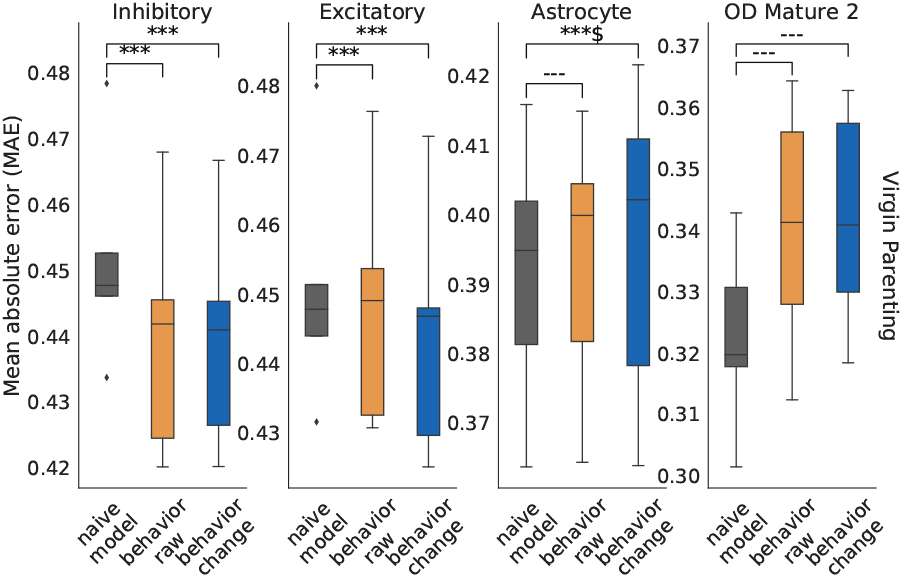
Comparison of naive and behavior models when the behavior is virgin parenting. Results are presented for the four major cell types. Naive model - predictions based on the naive model. Behavior raw - predictions based on learning from the raw values in the corresponding behavioral samples. Behavior change - predictions based on learning from the raw values in the corresponding behavioral samples subtracted by the predictions from the naive model. See Supplementary Methods for details. ***: p-value below 1e-3; **: p-value below 1e-2; *: p-value below 5e-2; ---: p-value larger than 5e-2; $: more than half of the CV groups show non-significant improvement when using behavioral models.

Given these differences, we next looked more closely at experts for which specific signalling genes have significant differences in their model coefficients when comparing the naive and behavioral models.

Figure 7 a) presents the coefficients MESSI learned from the raw values in naive, parenting, and virgin parenting animals for excitatory neurons. (For visualization purposes, we only show a subset of features and responses. See *Supplementary Methods* for how these were selected.) As can be seen, one of the top features with a much stronger impact for the parenting and virgin parenting models is oxytocin (Oxt), which has been shown to facilitate the onset and maintenance of maternal responsiveness in rodents’ brain (Rilling and Young, 2014). Further analysis of the MESSI results indicates that one of the experts (subtype 5 of parenting) has especially large weights for Oxt for a wide spectrum of response genes (Figure 7 a)). Examining the spatial location of the cells assigned to that subtype by MESSI indicates that these are likely oxytocin secreting magnocellular neurosecretory neurons located in the paraventricular nucleus (PVA) (Figure 7 b)).

**Fig. 7.**
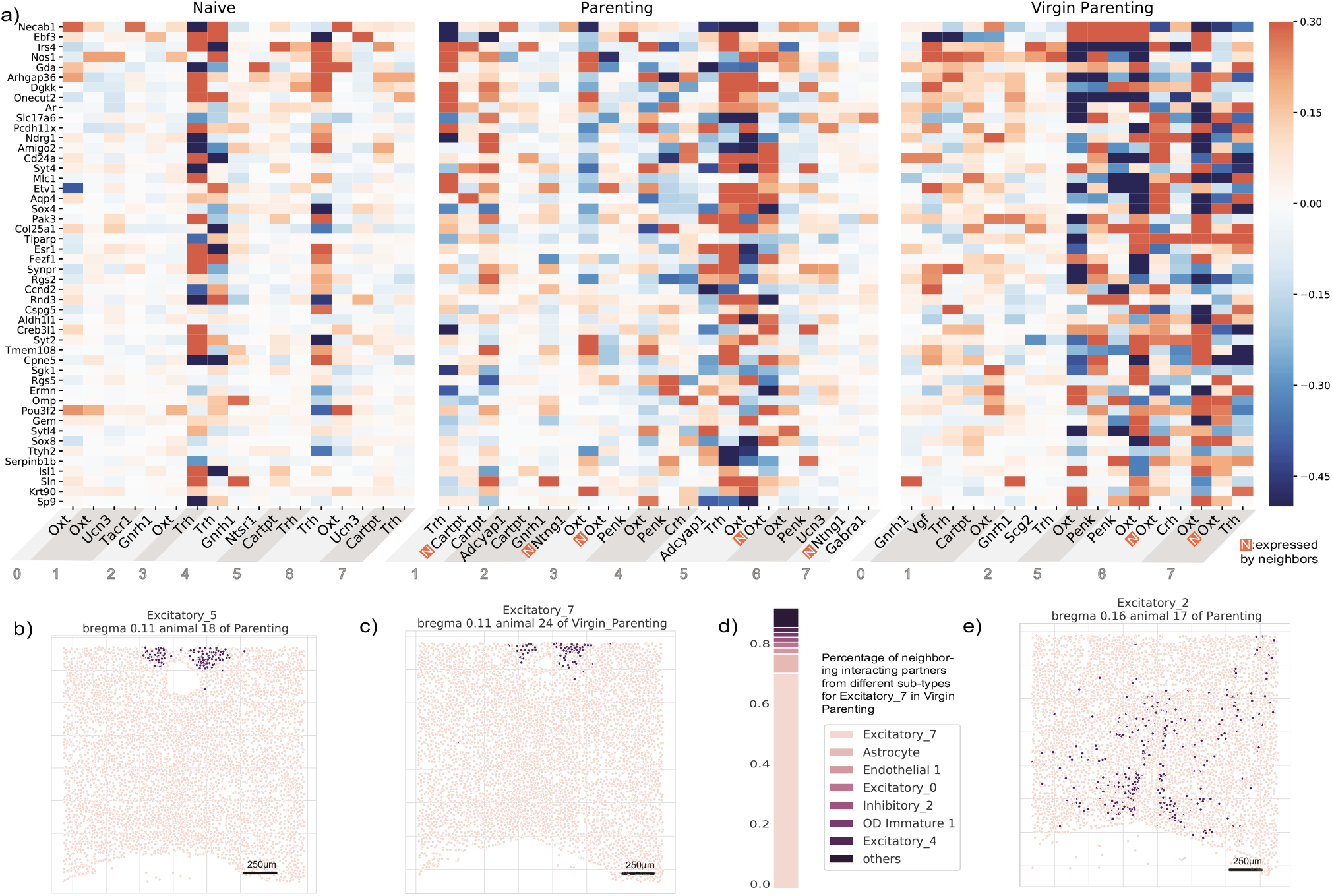
MESSI reveals changes in key signaling molecules and relevant signaling networks activated upon experience. Top a): Coefficients for top signaling molecules in a subset of the MESSI experts of excitatory cells for different conditions (X axis) for several response genes (Y axis). Note the large increase in Oxt coefficients between naive and parenting or virgin parenting models. See Supplementary Methods for the selection of top features. Bottom: Cells assigned to specific MESSI experts. b), c), e): spatial location of the cells on an example bregma; d): the proportion of interacting partners from each expert as indicated by the expression level of neighboring oxytocin

A subtype of oxytocin secreting cells located in the same region was also identified by MESSI for another experimental condition, virgin parenting (subtype 7 in Figure7 a) and c)). However, while we identified Oxt for both behaviors, the response genes influenced the most from Oxt for the virgin parenting model are different from those predicted to be impacted in the parenting animals. This may imply differences in mechanisms for Oxt secretion regulation. Indeed, for virgin parenting, we found that a major impact for Oxt arises from neighboring cells, whereas for parenting, its main impact is internal (Figure 7 a)). Unlike many other neuropeptides released primarily from axons, oxytocin is known to be exclusively released within the PVA from dendrites or somata and to act on nearby neighboring cells (Pow and Morris, 1989) further corroborating our hypothesis about the identity of this subtype. The differences in the signs for the coefficients of neighboring oxytocin and intracellular oxytocin may result from a negative feedback loop that impacts the production of oxytocin in nearby neurons.

Compared to the results from a clustering-based approach applied in the original paper (Moffitt *et al*., 2018), these two groups of cells (subtype 5 in parenting and subtype 7 in virgin parenting) correspond to cluster E-9 (80% overlapped). In line with their differential genes analysis results, we also found the genes Ntng1, Cck as top features in the classifiers determining if a cell belongs to these two subtypes (*Supplementary Figures* Figure S3). However, Oxt was not found to be an important gene in any of their clusters, likely because the amount of Oxt is very low in most excitatory cells.

Another neurohormone identified as differentially activated by MESSI is the gonadotropin-releasing hormone (GnRH), which is a major regulator of reproduction and may also play a role in maternal protection of offspring (Bayerl *et al*., 2019). GhRH1 was found to be significant for one of the other experts learned by MESSI (subtype 2 in parenting animals in Figure 7 a)). Another significant gene for this expert, pituitary adenylate cyclase-activating peptide (Adcyap1), has been shown to regulate gonadotropin subunits, both alone and in cooperation with GnRH in a pulse frequencydependent manner (Kanasaki *et al*., 2009). As shown in 7 e), this group of cells are mainly in the medial preoptic area (mPOA), as previously shown (Strauss and Barbieri, 2013). Interestingly, cFOS, an activation indicator of the consummatory aspects of maternal behaviors (Mattson and Morrell, 2005), is significantly enriched in this group of neurons (p-value < 1e-34 when compared to other neurons). We also found that the cocaine and amphetamine regulated transcript prepropeptide (Cartpt) has a larger impact in the parenting samples when compared to the naive or virgin parenting samples, as shown in Figure 7. CART immunoreactive neurons were found to be activated when postpartum females chose pup-over cocaine-associated environment in MPOA and other brain regions (Mattson and Morrell, 2005). These cells with Cartpt activated (expert 1 of parenting), which are primarily located in the bed nucleus of the stria terminalis (BNST) as shown in Figure S4 a), may also relate to pup exposure. A number of other potential subtypes with significant signaling genes, including proenkephalin (Penk), and Galanin (Gal), have also been identified using MESSI and are discussed in detail in *Supplementary Results*.

## 4 Discussion and future work

Recently developed methods for profiling spatial gene expressions at the single cell level open up new opportunities to study cell-cell communication. Most methods developed for the analysis of such data are unsupervised and focus on the clustering of cells or genes. To identify key genes involved in cell-cell interactions from spatial single cell data and allow quantitative evaluations, we developed MESSI, a novel predictive framework based on Mixture of Experts (MoE) and multi-task learning. MESSI predicts the expression of genes in a specific cell type using a combination of intra-cellular signaling genes and the expression of signaling genes in neighboring cells. Genes identified by MESSI as important for accurate predictions (based on their learned coefficients) are likely playing important roles in such interactions and can shed light on the impact of cell communication on the cell’s activity.

Testing MESSI on three recent datasets, we have shown that the method is able to achieve high accuracy, improving upon several previous methods used for gene expression prediction. Analysis of the resulting coefficients for the MERFISH hypothalamus dataset identifies Oxt and other signaling genes as important signaling genes in parenting or virgin parenting animals. Analysis of the subtypes (experts) in which these genes have a large impact indicates that they represent cells used to regulate parenting behavior. Unlike prior methods that are hard to interpret, MESSI can be directly used to identify key signaling genes and cell subtypes. Thus, both the fact that it is quantitatively better and the fact that it is more interpretable make it a better choice, we believe, for the analysis of spatial data.

MESSI’s high sensitivity in detecting signaling genes makes it a useful complement of the methods focused on identifying differentially expressed genes between cell types (subtypes). The expression of both Oxt and GnRH1 is very low in most excitatory neurons of the female parenting/virgin parenting animals in the MERFISH hypothalamus data. This likely explains why Oxt was not identified in the original paper and why the GnRH1 cluster contained only very few cells (Moffitt *et al*., 2018). In contrast, by looking at their impact on other genes in the network, MESSI was able to identify both as important signaling factors for the parenting/virgin parenting animals. Indeed, neuropeptides are known to be activated post-translationally, which can explain the relatively small changes in expression levels observed (Varro, 2007). Another possible reason is that oxytocin is secreted in pulses for lactation and parturition (Fink *et al*., 2012), and the dendritic release within PVA facilitates pulsatile secretion (Bealer *et al*., 2010), and so it is not universally high in all excitatory cells. GnRH is another classic example of a pulsatile secretion hormone whose transcription level oscillates in an episodic pattern (Choe *et al*., 2013). Thus, it is possible that methods that do not allow the analysis of gene-gene interactions in subtypes of cells (for example, those using clustering) miss such events, whereas MESSI can identify them using the different experts and their learned regression models.

While MESSI can accurately predict gene expression and is able to reveal the activity of key signaling genes, there are a number of ways in which it can be further improved. First, we noticed that some signaling genes are highly correlated. Such collinearity results in reduced importance assigned to each of the correlated genes such that the model may miss identifying some of the key signaling genes. A possible way to mitigate this problem is through the use of a more stringent regularization penalty. Another issue is the scaling to larger datasets with tens of thousands of profiled genes. Such data presents challenges related to computational efficiency and would require extensions to the current implementation that can account for large numbers of profiled genes.

MESSI was developed in Python and is available as an open-source package from https://github.com/doraadong/MESSI.

## Supporting information

Supplementary methods and results

## Funding

This work was partially funded by the National Institutes of Health (NIH) [grants 1R01GM122096 and OT2OD026682 to Z.B.J.].

## Notes

### Competing Interest Statement

The authors have declared no competing interest.

