## Supplementary methods and results for "Identifying signaling genes in spatial single cell expression data"

August 24, 2020

Dongshunyi Li [1], Jun Ding [1] and Ziv Bar-Joseph [1,2,*]

[1]Computational Biology Department and [2]Machine Learning Department, School of Computer Science, Carnegie Mellon University, Pittsburgh, PA 15213, United States.

[*]To whom correspondence should be addressed.

**3 Supplementary Figures & Tables** **15**

### 1 Supplementary Methods

#### 1.1 Training and applying MESSI to make predictions

##### 1.1.1 Derivation of optimization objectives

The *incomplete* log conditional likelihood is denoted as $\mathcal{L}(\Theta)$, where $\Theta = \{\theta_c, \theta_j \text{ for each expert}\}$ and $\theta_j = \{W_j, b_j, \Omega_j, \Sigma_j\}$. $\theta_c$ is the set of parameters used in the classifier. For example, for logistic regression these are the coefficients of the predictors of the classifier. The objective is to maximize $\mathcal{L}(\Theta) = \log p(Y \mid X; \Theta)$, by Jensen's inequality, we have

$$\begin{aligned}
\log p(Y \mid X; \Theta) &= \log \sum_{\mathbf{z}} p(\mathbf{z}, Y \mid X; \Theta) \\
&= \log \sum_{\mathbf{z}} q(\mathbf{z}) \frac{p(\mathbf{z}, Y \mid X; \Theta)}{q(\mathbf{z})} \\
&\geq \sum_{\mathbf{z}} q(\mathbf{z}) \log \frac{p(\mathbf{z}, Y \mid X; \Theta)}{q(\mathbf{z})} \\
&= \sum_{\mathbf{z}} q(\mathbf{z}) \log p(\mathbf{z}, Y \mid X; \Theta) - \sum_{\mathbf{z}} q(\mathbf{z}) \log q(\mathbf{z}). \qquad (1)
\end{aligned}$$

Define (1) as $Q(q, \Theta)$. Then the EM algorithm, as stated in Bilmes *et al.* (1998), iterates between

$$\begin{aligned}
E &: q^t = \underset{q}{\operatorname{argmax}}\, Q(q, \Theta^{t-1}) \\
M &: \Theta^t = \underset{\Theta}{\operatorname{argmax}}\, Q(q^t, \Theta),
\end{aligned}$$

where $t$ is the $t_{th}$ iteration. Given that the inequality above equals when $q(\mathbf{z}) = p(\mathbf{z} \mid Y, X; \Theta)$, we have $q^t(\mathbf{z}) = p(\mathbf{z} \mid Y, X; \Theta^{t-1})$ maximizes $Q(q, \Theta^{t-1})$. For M-step, $Q(q^t, \Theta) = \sum_{\mathbf{z}} q^t(\mathbf{z}) \log p(\mathbf{z}, Y \mid X; \Theta)$, given that the other term is constant with regarding to $\Theta^t$. We have

$$\begin{aligned}
E &: q^t(\mathbf{z}) = p(\mathbf{z} \mid Y, X; \Theta^{t-1}) \\
M &: \Theta^t = \underset{\Theta}{\operatorname{argmax}} \sum_{\mathbf{z}} p(\mathbf{z} \mid Y, X; \Theta^{t-1}) \log p(\mathbf{z}, Y \mid X; \Theta).
\end{aligned}$$

In E-step, for each cell $i$, we calculate

$$p(z_i = j \mid \mathbf{y_i}, \mathbf{x_i}; \Theta^{t-1}) = \frac{p(z_i = j \mid \mathbf{x}_i; \theta_c^{t-1}) p(\mathbf{y_i} \mid \mathbf{x_i}, z_i = j; \theta_j^{t-1})}{\sum_j^J p(z_i = j \mid \mathbf{x_i}; \theta_c^{t-1}) p(\mathbf{y_i} \mid \mathbf{x_i}, z_i = j; \theta_j^{t-1})}. \qquad (2)$$

In M-step, given the modularity of Mixture of Experts (MoE) models and the conditional independence among samples, we have

$$\begin{aligned}
\sum_{\mathbf{z}} p(\mathbf{z} \mid Y, X; \Theta^{t-1}) \log p(\mathbf{z}, Y \mid X; \Theta) &= \sum_i^N \sum_j^J p(z_i = j \mid \mathbf{y_i}, \mathbf{x_i}; \Theta^{t-1}) \log p(z_i = j, \mathbf{y_i} \mid \mathbf{x_i}; \Theta) \\
&= \sum_i^N \sum_j^J p(z_i = j \mid \mathbf{y_i}, \mathbf{x_i}; \Theta^{t-1}) \log p(z_i = j \mid \mathbf{x_i}; \theta_c) p(\mathbf{y_i} \mid \mathbf{x_i}, z_i = j; \theta_j) \\
&= \sum_i^N \sum_j^J p(z_i = j \mid \mathbf{y_i}, \mathbf{x_i}; \Theta^{t-1}) \log p(z_i = j \mid \mathbf{x_i}; \theta_c) + \sum_j^J Q_j(\theta_j),
\end{aligned}$$

where $Q_j(\theta_j)$ is the objective for the jth expert and $Q_j(\theta_j) = \sum_i^N p(z_i = j \mid \mathbf{y_i}, \mathbf{x_i}; \Theta^{t-1}) \log p(\mathbf{y_i} \mid \mathbf{x_i}, z_i = j; \theta_j)$. Denote $p(z_i = j \mid \mathbf{y_i}, \mathbf{x_i}; \Theta^{t-1})$ as $h_j(i)$. Then the above can be written as

$$\sum_{\mathbf{z}} p(\mathbf{z} \mid Y, X; \Theta^{t-1}) \log p(\mathbf{z}, Y \mid X; \Theta) = \sum_i^N \sum_j^J h_j(i) \log p(z_i = j \mid \mathbf{x_i}; \theta_c) + \sum_j^J Q_j(\theta_j). \tag{3}$$

That is, in M-step, the optimization task can be decomposed into optimizing a classifier with objective $Q_c(\theta_c) = \sum_i^N \sum_j^J h_j(i) \log p(z_i = j \mid \mathbf{x_i}; \theta_c)$ and optimizing each expert $j$ with objective $Q_j(\theta_j)$. Given we apply MROTS (Rai *et al.*, 2012) for each expert, by taking account of the prior specified in *Materials and Methods* and other regularizers, we have $Q_j(\theta_j)$ as

$$\begin{aligned} Q_j(\theta_j) = &\sum_i^N h_j(i) \log p(\mathbf{y_i} \mid \mathbf{x_i}, z_i = j; \theta_j) - \frac{\lambda}{2}\mathrm{tr}(W_j W_j^T) - \frac{\lambda_1}{2}\mathrm{tr}(W_j \Sigma_j^{-1} W_j^T) \\ &+ \frac{D}{2}\log|\Sigma_j^{-1}| - \frac{\lambda_2}{2}\mathrm{tr}(\Omega_j^{-1}) - \frac{\lambda_3}{2}\mathrm{tr}(\Sigma_j^{-1}). \end{aligned} \tag{4}$$

Note that if we assume sparsity of $\Omega^{-1}$ and $\Sigma^{-1}$, then the regularizers $\mathrm{tr}(\Omega_j^{-1})$ and $\mathrm{tr}(\Sigma_j^{-1})$ are replaced by the $l_1$ norm regularizers $\|\Omega_j^{-1}\|_1$ and $\|\Sigma_j^{-1}\|_1$.

##### 1.1.2 Derivation of updates for experts

As suggested by Rai *et al.* (2012) and Cai *et al.* (2014), MROTS models are estimated by alternating minimization. That is, we update a parameter at a time sequentially assuming the other parameters unchanged. Here we derive the updates for each parameter of a MROTS expert $j$ at a single iteration $t$:

1. Update $W_j$, assuming $\mathbf{b_j}, \Omega_j, \Sigma_j$ unchanged

$$\begin{aligned} \frac{\partial Q_j}{\partial W_j} &= \frac{\partial \sum_i^N h_j(i) \log p(\mathbf{y_i} \mid \mathbf{x_i}, z_i = j; \theta_j) - \frac{1}{2}\lambda \mathrm{tr}(W_j W_j^T) - \frac{1}{2}\lambda_1 \mathrm{tr}(W_j \Sigma_j^{-1} W_j^T)}{\partial W_j} \\ &= \frac{\partial \sum_i^N h_j(i) \log p(\mathbf{y_i} \mid \mathbf{x_i}, z_i = j; \theta_j)}{\partial W_j} - \lambda W_j - \lambda_1 W_j \Sigma_j^{-1} \\ &= \sum_i^N h_j(i) \frac{\partial p(\mathbf{y_i} \mid \mathbf{x_i}, z_i = j; \theta_j)}{\partial W_j} \frac{1}{p(\mathbf{y_i} \mid \mathbf{x_i}, z_i = j; \theta_j)} - \lambda W_j - \lambda_1 W_j \Sigma_j^{-1} \\ &= \sum_i^N h_j(i) \mathbf{x_i} (\mathbf{y_i} - W_j^T \mathbf{x_i} - \mathbf{b_j})^T \Omega_j^{-1} - \lambda W_j - \lambda_1 W_j \Sigma_j^{-1} \end{aligned}$$

Set $H_j$ as a diagonal matrix with diagonal terms $d_j(i) = h_j(i)$. By setting $\frac{\partial Q_j}{\partial W_j} = 0$, we have

$$\big(\Omega_j^{-1} \otimes X^T H_j X + (\lambda_1 \Sigma_j^{-1} + \lambda I_K) \otimes I_D\big)\mathrm{vec}(W_j) = \mathrm{vec}(X^T H_j (Y - \mathbf{1} b_j^T) \Omega_j^{-1}), \tag{5}$$

where $\mathbf{1}$ is a $N \times 1$ vector and vec(A) is the vectorized form of a matrix $A$. Then we solve the linear system above and obtain the update for $W_j$. This can be regarded as a weighted generalization of the original derivation in Rai *et al.* (2012) and Cai *et al.* (2014). Note that Cai *et al.* (2014) proposed an alternative solution when the number of tasks $K$ is big by applying singular value decomposition (SVD) instead of solving a linear system. Given that we only have a moderate number of tasks $K$, we now obtain the updates by solving the linear system. We will consider solving by SVD when the task number increases substantially in the future.

2. Update $\mathbf{b_j}$, assuming $W_j, \Omega_j, \Sigma_j$ unchanged

$$\begin{aligned}\frac{\partial Q_j}{\partial \mathbf{b_j}} &= \sum_i^N h_j(i) \frac{\partial p(\mathbf{y_i} \mid \mathbf{x_i}, z_i = j; \theta_j)}{\partial \mathbf{b_j}} \frac{1}{p(\mathbf{y_i} \mid \mathbf{x_i}, z_i = j; \theta_j)} \\ &= \sum_i^N h_j(i) \Omega_j^{-1} (\mathbf{y_i} - W_j^T \mathbf{x_i}) - \sum_i^N h_j(i) \Omega_j^{-1} \mathbf{b_j}\end{aligned}$$

Denote $\mathbf{h_j}$ as a $N \times 1$ vector with element as $h_j(i)$. By setting $\frac{\partial Q_j}{\partial \mathbf{b_j}} = 0$, we have

$$\begin{aligned}\mathbf{b_j} &= \frac{\sum_i^N h_j(i)(\mathbf{y_i} - W_j^T \mathbf{x_i})}{\sum_i^N h_j(i)} \\ &= \frac{1}{\sum_i^N h_j(i)} (Y - XW_j)^T \mathbf{h_j}. \end{aligned} \tag{6}$$

Again, it can be regarded as a weighted generalization of the original derivation in Rai *et al.* (2012) and Cai *et al.* (2014).

3. Update $\Omega_j$, assuming $\mathbf{b_j}, W_j, \Sigma_j$ unchanged

$$\begin{aligned}\frac{\partial Q_j}{\partial \Omega_j^{-1}} &= \frac{\partial \sum_i^N h_j(i) \log p(\mathbf{y_i} \mid \mathbf{x_i}, z_i = j; \theta_j) - \frac{1}{2}\lambda_2 \mathrm{tr}(\Omega_j^{-1})}{\partial \Omega_j^{-1}} \\ &= -\frac{1}{2}\Big(\sum_i^N h_j(i) \frac{\partial \mathrm{tr}((\mathbf{y_i} - W_j^T \mathbf{x_i} - \mathbf{b_j})^T \Omega_j^{-1} (\mathbf{y_i} - W_j^T \mathbf{x_i} - \mathbf{b_j}))}{\partial \Omega_j^{-1}} - \sum_i^N h_j(i) \frac{\partial \log(\Omega_j^{-1})}{\partial \Omega_j^{-1}} + \lambda_2 \frac{\partial \mathrm{tr}(\Omega_j^{-1})}{\partial \Omega_j^{-1}}\Big) \\ &= -\frac{1}{2}\Big(\sum_i^N h_j(i)(\mathbf{y_i} - W_j^T \mathbf{x_i} - \mathbf{b_j})(\mathbf{y_i} - W_j^T \mathbf{x_i} - \mathbf{b_j})^T - \sum_i^N h_j(i)\Omega_j + \lambda_2 I_K\Big)\end{aligned}$$

By setting $\frac{\partial Q_j}{\partial \Omega_j^{-1}} = 0$, we have

$$\Omega_j = \frac{(Y - XW_j - \mathbf{1}b_j^T)^T H_j (Y - XW_j - \mathbf{1}b_j^T) + \lambda_2 I_K}{\sum_i^N h_j(i)}. \tag{7}$$

Again, it can be regarded as a weighted generalization of the original derivation in Rai *et al.* (2012) and Cai *et al.* (2014). As suggested by Rai *et al.* (2012), if we assume sparsity for $\Omega^{-1}$, then we can instead solve a graphical lasso problem for the updates. See Rai *et al.* (2012) for details.

4. Update $\Sigma_j$, assuming $\mathbf{b_j}, W_j, \Omega_j$ unchanged
Given updating $\Sigma_j$ is not directly depending on the data $Y$ and $X$, the rule stays the same as in Rai *et al.* (2012) and Cai *et al.* (2014). As suggested by Rai *et al.* (2012), if we assume sparsity for $\Sigma^{-1}$, then we can instead solve a graphical lasso problem for the updates. Please refer to the original papers of MROTS for the details. For the update, we have

$$\Sigma_j = \frac{\lambda_1 W_j^T W_j + \lambda_3 I_K}{D}. \tag{8}$$

##### 1.1.3 Updates for the classifier/gate

The objective for the classifier is $Q_c(\theta_c) = \sum_i^N \sum_j^J h_j(i) \log p(z_i = j \mid \mathbf{x_i}; \theta_c)$. As suggested by Jordan and Jacobs (1994), a popular choice for the classifier is logistic regression. Then this logistic regression with proportions (as compared to labels) as outputs can be learned by Iterative Reweighted Least Squares (IRLS). However, in practice, we found that this approach did not perform as well as training based on labels in our particular application. Thus, we tweak the learning objective slightly by replacing the proportions $h_j(i)$ with labels. Given that we assume each cell $i$ is generated by a single expert, we assign cell $i$ with label $j$ if $h_j(i) = \max\{h_1(i), ..., h_s(i), ..., h_J(i)\}$. That is, we put all probability mass on $j$ that is most likely. Under this transformation, instead of using $h_j(i)$ in the objective $Q_c(\theta_c)$, we use an indicator defined in the following:

$$\mathbb{1}(z_i = j; \Theta^{t-1}) = \begin{cases} 1, & \text{if } j = \text{argmax}_s\{h_1(i), ..., h_s(i), ..., h_J(i)\} \\ 0, & \text{otherwise.} \end{cases} \tag{9}$$

And we have the tweaked objective $Q'_c(\theta_c)$ as

$$Q'_c(\theta_c) = \sum_i^N \sum_j^J \mathbb{1}(z_i = j; \Theta^{t-1}) \log p(z_i = j \mid \mathbf{x_i}; \theta_c). \tag{10}$$

This transforms the classifier training problem into a standard label-based logistic regression. We apply the implementation by the open-source software *scikit-learn* (Pedregosa *et al.*, 2011) for the classifier.

##### 1.1.4 Updates for experts when assuming "hard" weights

The same tweak described above can also be applied to the parameter updates for experts. We can also use the indicator as defined in (9) in place of the "soft" $h_j(i)$. The other parts of the equations stay the same. In practice, we found this tweak gave better prediction accuracy for some cell types (of which subtypes might be more different from each other).

##### 1.1.5 Initialization

The EM algorithm is known to be sensitive to initialization. We found initializing the weights $h_j(i)$ for each cell $i$ and each expert $j$ by hierarchical clustering gave a good performance in practice. We apply the agglomerative clustering with the "ward" linkage implemented by the open-source software *scikit-learn* (Pedregosa *et al.*, 2011) for this purpose. For the parameters in expert $j$, we initialize $W_j$ and $b_j$ with 0. Note that this only applies to the first iteration. For the iterations after the first one, the estimates obtained in the previous iteration will be used as the initialization (so-called "warm-start").

##### 1.1.6 Stopping criterion

The M-step objective, as defined in equation (3), is normally applied as the stopping criterion. In practice, we use the error, that is, the cross-entropy error from the classifier and the mean squared error from the experts weighted by $h_j(i)$, as the stopping criterion.

##### 1.1.7 Training algorithm

See Algorithm 1.

**Algorithm 1** Training algorithm for MESSI models

**Input:** $X \in \mathbb{R}^{N \times D}$, $Y \in \mathbb{R}^{N \times K}$; $J$ as number of experts, $\lambda$, $\lambda_1$, $\lambda_2$, $\lambda_3$ for MROTS
**Output:** $\theta_c$ for the classifier; $\theta_j = \{W_j, \mathbf{b_j}, \Omega_j, \Sigma_j\}$ for each expert $j$

```
Initialize h_j(i) for each cell i and each expert j by hierarchical clustering
t = 0
while not converged do
    for j = 1, 2, .., J do
        W_j ← solution to the linear system defined in (5)
        b_j ← equation (6)
        Ω_j ← equation (7)
        Σ_j ← equation (8)
    end for
    θ_c ← train a logistic classifier with objective defined as equation (10)
    h_j(i) ← equation (2)
    t = t + 1
end while
```

###### 1.1.8 Predictions

For a new sample $\mathbf{x} \in \mathbb{R}^D$, the prediction $\hat{\mathbf{y}} \in \mathbb{R}^K$ equals $\mathbb{E}(\mathbf{y} \mid \mathbf{x}) = \sum_j^J \mathbb{E}(\mathbf{y} \mid \mathbf{x}, z = j) p(z = j \mid \mathbf{x})$. $\mathbb{E}(\mathbf{y} \mid \mathbf{x}, z = j)$ is the predicted value from expert $j$ and is simply given by $W_j^T x + b_j$. $p(z = j \mid \mathbf{x})$ is the output from the classifier given input vector $\mathbf{x}$. Thus, the prediction $\hat{\mathbf{y}}$ can be viewed as a weighted average over the predictions made by different experts.

###### 1.1.9 Nested cross-validation (CV)

Following the introduction of the main idea in *Materials and Methods*, here we describe in detail how we conduct a grid search for the hyperparameters in the inner loop. Given that the datasets we use all have very limited number of animals/replicates (4 for the MERFISH hypothalamus datasets, 3 for the other two datasets), and we leave one animal/replicate out for evaluation purpose in the outer loop, we have very few of them left for the inner loop. If we still split the data only based on the animal/replicates, then the size of the validation set would be similar to the training set. Thus, we decide to split the cells also based on their spatial proximity. For example, for the MERFISH hypothalamus dataset where a third spatial dimension, bregma, is also measured for each cell, we split the data by the bregma as well as the animal. The cells that are closed to each other on the 2D plane corresponding to a slide (bregma) will be kept together and their spatial relationship will be preserved. For the other 2 datasets that are only measured for 2 spatial dimensions, we instead employ a sliding window approach and divide a single replicate into several regions based on spatial proximity.

For each iteration of the inner loop, we randomly split the original training set into a new training set and a validation set based on the above criteria at a user-defined validation/training ratio (the default is 0.2). The number of iterations, equivalently the number of training and validation pairs, is also user-specified (the default is 5).

For each hyperparameter, we specify the possible values it can take before conducting the grid search. For example, for the number of experts, by default we have 4 different candidate values to loop through for all datasets.

#### 1.2 Data processing

##### 1.2.1 Selection of ligand/receptors for features

Out of all profiled genes, we choose the profiled genes that are labeled as ligand or receptor according to a database of interacting ligands and receptors (Ramilowski *et al.*, 2015) and use all of them for features. This database consists of 708 ligands, and 691 receptors with 2,557 known interactions between ligands and receptors (Ramilowski *et al.*, 2015). By manual curation, we found a few genes not included in this database but are likely ligand or receptor expressed in the brain (for example, Gabra1). We also use them as ligand/receptors. See Table S1 in *Supplementary Figures & Tables* for this list of genes.

For the MERFISH hypothalamus dataset, 71 genes are indicated by the list from Ramilowski *et al.* (2015) and Table S1 as ligands or receptors. For the STARmap data, 39 out of 166 genes profiled are indicated as ligand/receptors. The MERFISH U-2 OS paper (Xia *et al.*, 2019) identified a set of 742 DE genes of clusters, of which 49 are annotated as ligand/receptors.

##### 1.2.2 Selection of genes as response variables

For the MERFISH hypothalamus dataset, we use the full list of profiled genes in the modeling. Except for those used as features (ligand/receptors), we have 84 genes used as response variables. We also use the full list of profiled genes for the STARmap data, resulting in 127 genes as response variables. For the MERFISH U-2 OS dataset, we use those that are identified as DE (differentially expressed) genes in different clusters constructed by the original study (Xia *et al.*, 2019). We select the union of top 10, 20, 50, 100 and all DE genes of each cluster, except for those annotated as ligand/receptors, resulting in the number of response variables as 38, 74, 181, 364 and 693, respectively.

##### 1.2.3 Construction of neighborhood graphs

Given the coordinates of cells' centroid, we construct neighborhood graphs by applying Delaunay triangulation with certain distance cutoff (for example, 100 micrometers for the MERFISH hypothalamus dataset), as illustrated at the left-hand side of Figure 1. We then find the first-order neighbors for every node (cell) in the graph and regard them as the neighbors for each cell. Given the datasets we used are 2D, without knowing the (continuous) spatial information in the third dimension, we only find neighbors within a single bregma (MERFISH hypothalamus data)/batch (MERFISH U-2 OS data)/sample (STARmap mPFC data).

##### 1.2.4 Preparation for features and responses

We consider the following features: the expression of ligands and receptors in cell $i$, the expression of ligands in cell $i$'s neighbors, cell $i$'s spatial coordinates (location within the image/tissue) and cell types of its neighbors (we used the cell type annotations from the original studies). We take log transformation of all genes for both features and responses. We take the expression values of ligands and receptors expressed by the cell itself as the intra-cellular features. For an inter-cellular feature (a ligand), we take the sum over the expression of the ligand in all its neighbors as the feature. For the MERFISH hypothalamus data, we use the bregma, x, and y coordinates of the cell centroid as the spatial position feature (for baseline models only "bregma" is used as stated in Section 1.3). The other two datasets do not provide the third dimension information, and thus we only use the x and y coordinates of the centroids of cells. As for the neighboring cell types, we count, for each cell type, how many times it appears in the cell's neighborhood and use it as the feature. Finally, we concatenate the different groups of features together and standardize all features.

###### 1.2.5 Features applied to different modules of MESSI

Out of the different types of features, we only apply the spatial position, and the neighboring cell type features to the classifier. In other words, the expert only models the relationship between the response genes and the intra- & inter- cellular expression of signaling molecules.

#### 1.3 Training baseline and benchmarking models

To construct the features for the baseline model, we apply the one-hot encoding for cell types and concatenate it with spatial coordinates (up to 2 dimensions depending on the specific dataset). Given that the slides (with x, y coordinates) from the same animal in the MERFISH hypothalamus dataset are not aligned and based on Figure S11, the performance with (x, y) coordinates is similar as that without, we decide not to include (x, y) coordinates in the baseline model for the MERFISH hypothalamus dataset. We use all coordinates available for the other datasets for the baseline model. We also generate second-degree polynomial features allowing interactions and use lasso by the open-source software *scikit-learn* (Pedregosa *et al.*, 2011) for training. We train all cell types together. We will describe how the penalty value is selected in Section 1.4.1.

The benchmarking methods have all been applied to gene expression prediction using a subset of feature genes (Chen *et al.*, 2016; Li *et al.*, 2019). Among them, Chen *et al.* (2016) described that researchers in LINCS program (http://www.lincsproject.org/) built a linear regression model for each response gene assuming response genes are conditionally independent. Li *et al.* (2019) applied the XGBoost (Chen and Guestrin, 2016) algorithm and also modeled different response genes separately. Chen *et al.* (2016) learned multi-layer perceptron (MLP) models with multi-node output and thus enabled weights sharing across response genes. We also included as a comparison, a MLP model with single-node output (thus without sharing among response genes). These methods are all based on popular machine learning models that have been implemented and optimized by popular open-source softwares. For linear regression, multi-node output MLP and single-node output MLP, we apply the implementation in *scikit-learn* (Pedregosa *et al.*, 2011). And for Li *et al.* (2019), we apply the XGBoost implementation from the original publication (Chen and Guestrin, 2016).

Among the benchmarking methods, MLP is known to be sensitive to the number of hidden nodes and the number of hidden layers. From preliminary results (data not shown), we found MLP with 1 hidden layer performed best in this task. Thus, we use 1 hidden layer for both MLP with single-node and multi-node output. We adjust the number of hidden nodes according to the sample size. For example, for the MERFISH hypothalamus data, we use 8 hidden nodes for the inhibitory cell type (55K sample size), 4 for astrocyte (17K sample size), and 2 for endothelial 1 (6K sample size) and the others with smaller sample sizes.

#### 1.4 Model evaluation & analysis

###### 1.4.1 Cross-validation strategies for the baseline and the benchmarking models

For the baseline model, to select the best value for the alpha (penalty) parameter as well as evaluate the performance, we employ the same nested CV strategy as we designed for MESSI (See Section 1.1.9 and Section 2.4 in *Materials and Methods*). The best alpha values, out of 10 values ranging from $10^{-4}$ to $10^{-0.5}$, selected for different cell types and datasets are listed in Table S2. For the benchmarking models, in each iteration we leave out one of the animals/replicates and learn the models using the remaining animals/replicates, which is equivalent to the outer loop of the nested CV strategy we employ for MESSI and the baseline model.

###### 1.4.2 Calculation of the error metric

We use mean absolute error (MAE) as the error metric. MAE is calculated as the arithmetic mean of the absolute differences in log-expression between the prediction and the true value. We first calculate the absolute values of the differences for every gene $k$ in every cell $i$ and obtain $|\log(\frac{y_{ik}}{\hat{y}_{ik}})|$, where $y$ denotes the true raw counts and $\hat{y}$ denotes the predicted raw counts. We then take the arithmetic mean over all response genes for each cell and finally take the arithmetic mean over all cells and obtain the MAE.

###### 1.4.3 Boxplots for CV evaluation results

For each cross-validation (CV) run (or equivalently the outer iteration of the nested CV strategy), we hold out one animal/replicate for testing. For each CV run, we calculate an MAE on the held-out animal/replicate. Unless specifically specified, the boxplots describe the MAEs from all (outer) CV runs.

###### 1.4.4 Significance test for comparing CV results

For two sets of CV results (each corresponding to a model or a condition), a one-sided Wilcoxon signed-rank test between the errors per cell (absolute error averaged over response genes only) obtained from the 2 models/conditions is performed for each test animal/replicate. The individual p-values are combined by Fisher's method (Mosteller and Fisher, 1948), which outputs a combined p-value. Unless specifically specified, the p-values shown on boxplots for CV evaluation are all combined p-values obtained through the approach described above. If the p-value is lower than 1e-3, then "***" will be shown on the plot. Likewise, "**" is for p-values lower than 1e-2 and "*" for 5e-2.

By the definition of Fisher's method, a small combined p-value (for example, lower than 5e-2) suggests that it is likely not all the individual null hypotheses are true. Thus, the combined p-value may be small, while 3 out of 4 CV runs show non-significant improvements by a certain model/condition. For visualization purposes, we marked the cases where the combined p-value is significant while more than half of the CV runs resulting in MAEs larger than the model/condition in comparison by "$" on the boxplots.

Unless specifically specified, the strategy described above is applied to every pairwise comparison of CV results in our study, for example, when testing if adding neighborhood information improves performance, when testing if MESSI outperforms another model and when testing if behavioral models outperform the naive model on predicting behavioral samples.

###### 1.4.5 Down-sampling experiment

For the MERFISH hypothalamus data, instead of combining all bregmas of a single animal together, we train the models using only samples from a single bregma at a time. This left us with 35 CV groups with training sample sizes ranging from 0.7K to 6K. We apply the same cross-validation (CV) strategies for MESSI, the baseline model, and the other benchmarking models, as described in Section 1.1.9 and 1.4.1. Same as what we do for the full data model, we do not include the x, y coordinates in the baseline model.

###### 1.4.6 Behavior change samples preparation

We first apply models learned from naive animals to make predictions for the behavioral samples. Then we subtract the predictions based on the model learned from naive animals from the behavioral samples' raw expression values. The resulting values can be regarded as the "change" of expression from being naive to displaying certain behaviors. Then we train models using these "change" of expression values.

###### 1.4.7 Comparison between naive and behavioral models on predicting behavioral samples

For the comparison between naive and behavioral (parenting or virgin parenting) models on predicting behavioral samples of the MERFISH hypothalamus dataset, we train the naive models using all available naive animals (in total 4) and use the models to make predictions on each behavioral animal. For the behavioral models, we conduct cross-validations (CV) for each behavior separately. We leave out one behavioral animal for testing and use the rest to learn the model. For both naive and behavioral models trained for this purpose, we do not apply the nested CV strategy. We instead fix the number of experts as shown in Table S4.

###### 1.4.8 Number of experts used for inference purposes

We also train models on all animals of a certain behavior with a fixed number of experts for inference purposes (for example, analyzing the coefficients and subtypes). See Table S5 for the number of experts we used.

###### 1.4.9 Selecting top features for visualizing coefficients

We select a subset of features as "top" features based on the learned coefficients. For each expert of each cell type, we select the features whose weight for any of the responses is above the 99.9% quantile of all weights in the same expert. Among the features selected for the excitatory cells shown on Figure 7, a few of them are not the "top" features but included as references. These include Cartpt_n in expert 1 of parenting, Oxt_n in expert 3 of parenting, Crh in expert 4 of parenting, Oxt_n and Adcyap1 in expert 5 of parenting, Penk and Oxt_n in expert 6 of virgin parenting (the suffix "_n" denotes features expressed from neighbors). For visualization, we use the original coefficients for heatmaps with cutoffs of -0.5 and 0.3 for the anchor points of the color map.

###### 1.4.10 cFOS enrichment p-value calculation

We do not include cFOS gene in modeling and use it only for validation purposes. We then apply similar procedures in Moffitt *et al.* (2018) for cFOS validation. Briefly, we first label cells as "cFOS enriched" if their cFOS expression is above 2 standard deviations of the mean of cFOS expression over all neurons in the same behavioral group. And we can obtain the null ratio, which is the number of cFOS enriched cells over all neurons in the same behavioral group. Then the enrichment p-value for a given group of cells is calculated by assuming a binomial distribution with the null distribution parameter equal the null ratio. The number of successes equals the number of cFOS enriched cells, and the number of total cases equals the total number of cells in the group.

###### 1.4.11 Proportion of interacting neighboring partners

For a given neighboring feature (that is, the feature from summing the expression of a certain ligand over all neighboring cells) and for each cell, we can identify which neighboring cells express the given ligand. These cells are regarded as contributing to the neighboring feature. We then calculate the proportion of a neighboring ligand from each cell subtypes (experts) learned from the MESSI models. The subtype that contributes the most is labeled as the "top contributor". Then we count how many times, in the whole dataset, that a cell subtype being the "top contributor". The output is then the fraction of times across all observations that a certain subtype being the "top contributor".

### 2 Supplementary Results

#### 2.1 Performance and inference when using different numbers of responses

We tested MESSI using different responses sets with various sizes. For this, we used the MERFISH U-2 OS dataset to vary the number of top DE genes selected as response genes between 38 and 693 (See Section 1.2.2 for selection details). We assume covariance structures for the experts except when we have 693 responses where the current implementation of MROTS requires large memory (an alternative implementation that solves the memory issue while still assuming covariance is discussed in Section 1.1.2). As Figure S7 shows, MESSI greatly improved upon the baseline model and performed better than all benchmarking models for all responses gene sets. We note that the MAE dropped when using more response genes, regardless of the methods. Given that the MAE, by definition, as stated in Section 1.4.2, is an average over all response genes, it is likely that the newly added genes are easier to predict. This was indeed the case as we show in Figure S8 a), where we plot the MAE of new genes added each time when more DE genes were included. The newly added genes, which are less differentially expressed than the ones already included, also have lower dispersion (variance/mean), shown by Figure S8 c).

Given that our goal is not the analysis of response genes but rather the identification of important signaling genes and cell subtypes, we checked how much impact the newly added response genes had on the identification of signaling genes. For each model using a specific set of response genes, we selected the top features based on learned coefficients, as described in Section 1.4.9. As shown in Figure S9, we observed that all top features selected when using a smaller number of response genes were also selected when the number of response genes increased.

#### 2.2 Adding neighborhood information overfitted MESSI for small datasets

In line with our down-sampling results showing that MESSI subjected to overfitting when the training sample size is small, we found that for small cell types (such as 2K OD Mautre 1 in Figure S20) of the MERFISH hypothalamus dataset and excitatory cells in the STARmap data (around 1.5K training size; Figure S22), MESSI performed worse when neighborhood information was included. In contrast, other models, such as XGBoost, still benefited from adding neighborhood information. However, when the training sample size is around 0.7K (for example, the MERFISH U-2 OS cells as shown in Figure S10), all models when including neighborhood information performed worse and were likely overfitted.

#### 2.3 Improvement from prediction performance of the behavior model unlikely due to sample size change

As stated in *Results*, we compared the performance of models learned using the naive or behavioral training samples on the task of predicting expression values for unseen behavioral samples. When using this approach for the MERFISH hypothalamus data, we observed an improvement in the accuracy from the behavioral models compared to the naive models for inhibitory and excitatory cell types. Given that we found a lack of data may lead to overfitting of the models, one possible reason for the improvement of performance by training using behavioral samples is the increased number of training samples. However, this is unlikely because that the training sample size of the behavioral samples is roughly 3 times fewer than the naive samples (for example, 20K inhibitory cells in parenting versus 66K in naive (when using all available animals) and 8.4K excitatory cells in parenting versus 31K in naive (when using all available animals)). Thus, the improvement in prediction ability is not likely due to the sample size differences.

#### 2.4 Classifier weights correspond to DE genes

The multi-class logistic regression model learns a set of coefficients for each expert. For a set of coefficients corresponding to a particular expert, we rank the features by coefficients. Each cell assigned to this expert also has a cluster label from Moffitt *et al.* (2018). We can compare the differential expressed genes assigned to these clusters by Moffitt *et al.* (2018) and the genes with the largest coefficients for this expert. Interestingly, some top features of some experts are also the DE genes of the corresponding clusters. For example, both excitatory expert 5 in parenting and expert 7 in virgin parenting correspond to E-9 cluster (by "correspond", we mean the majority of cells assigned to these experts also assigned with "E-9" label), which has Ntng1, Cck as its DE genes. Also, expert 1 in virgin parenting corresponds to E-5 with Penk as the DE gene. See Figure S3 for the representative experts.

#### 2.5 Identity of the cells assigned to excitatory expert 5 in parenting/expert 7 in virgin parenting

Most cells from either excitatory expert 5 in parenting or expert 7 in virgin parenting are assigned with E-9 cluster from Moffitt *et al.* (2018). According to Moffitt *et al.* (2018), E-9 cluster mostly locate in Paraventricular nuclei of thalamus (Pa/PVT), not Paraventricular nucleus of hypothalamus (PVA/PVN/PVH), even though they used PVA as the abbreviation. However, according to the anatomic reference in Allen Brain Atlas (sagittal sections of mouse p56) (Lein *et al.*, 2007), PVA locates closer to the medial preoptic area (mPOA) than Pa. Given that we found oxytocin and neighboring oxytocin very important for these 2 experts, we think it is likely that there exists oxytocin secreting magnocellular cells among these subgroups of cells.

It is also likely that these subgroups represent a hybrid of cell populations. We observed from Figure 7 in *Results* that for both excitatory expert 5 in parenting and expert 7 in virgin parenting, Trh is also very important. It was discovered that parvocellular Trh neurons are also in the magnocellular division of PVA (Kádár *et al.*, 2010) and are intermingled with magnocellular oxytocin cells. Thus, it is likely that these subgroups defined by the 2 experts represent a mixture of cells from both magnocellular Oxt and parvocellular Trh cells. The size of subgroups corresponding to these 2 experts is very small (about 0.2K cells), and we expect that MESSI can better separate them if more samples are available.

#### 2.6 Oxytocin receptor expressed higher in excitatory expert 5 in parenting/expert 7 in virgin parenting

We also found that the expression of oxytocin receptor (Oxtr) is significantly higher in subtype 5 in parenting and subtype 7 in virgin parenting, when compared to all other excitatory cells in the same behavior group as shown in Table S6.

#### 2.7 Other interesting experts

##### 2.7.1 Excitatory cells

We also found expert 4 and expert 6 from the parenting model both with oxytocin (Oxt) and proenkephalin (Penk) significantly affecting multiple response genes. As shown in Figure S4 b) and c), both groups of cells are located primarily in the anteroventral periventricular nucleus (AvPe / AVPV). Interestingly, we found in expert 6, Penk strongly associates with CAMP responsive element binding protein 3 like 1 (CREB3l1) (See Figure 7 in *Results*). Penk is a target of CREB, and it has been shown that phosphorylated CREB (pCREB) in AvPe may mediate the effect of steroid hormones on regulating Penk expression (Gu *et al.*, 1996).

###### 2.7.2 Inhibitory cells

Galanin (Gal) expressing inhibitory neurons in MPOA has been shown as a coordinating center for maternal behaviors (Wu *et al.*, 2014). We found expert 4 in parenting animals with Gal as the sole significant signaling molecule. As shown in Figure S5 a), this group of cells locates in both MPOA and ventrolateral preoptic nucleus (VLPO) (See Figure S5 b)), where galanin (Gal) neurons were found involved in sleep regulation (Kroeger *et al.*, 2018). Thus, this subtype is likely a mixture of 2 functionally different groups, both secreting galanin. As shown in Figure S5 d), these cells were mostly predicted to be subtype 2 by the naive model where Gal is also the main signaling molecule and with a similar profile of associations with response genes. Together with the finding that cFOS is not significantly enriched, this evidence indicate that this signaling subtype represents mostly Gal neurons that are not particularly activated in parenting behavior.

In contrast, we did find the subtype (expert) 7 in parenting, with very significant cFOS enrichment p-value ($p < 1e - 36$) locating in MPOA, as shown in Figure S5 c). The selected signaling molecule, somatostatin (Sst), a major regulator for growth hormone and many gastrointestinal hormones, implies a possible cross-talk between the neuronal circuits in feeding and that in parenting. Indeed, as shown in Figure S5 a), Sst in subtype 7 exerts a strong negative impact on insulin receptor substrate 4 (Irs4), which serves as an interface for multiple growth factor receptors, including receptors for Igf1. Igf1 in MPOA has been shown to be associated with maternal behavior regulation (Lékó *et al.*, 2017). Comparing to the clustering results by Moffitt *et al.* (2018), the major cell population (76%) of this expert corresponds to different clusters from Moffitt *et al.* (2018) that are all mapped with a single scRNA-seq cluster with Gal as the DE gene. Gal was selected by MESSI as one of the top features of the classifier assigning cells to this expert (Figure S3 expert 7 in parenting inhibitory). The reason why Gal was not selected as a significant signaling gene in the expert is likely due to the strong collinearity between Gal and other features such as bombesin receptor subtype 3 (Brs3) and calcitonin receptor (Calcr), as shown in Figure S6. In comparison, Sst does not have obvious collinearity with any other ligand, and thus we can easily distinguish its influence from the others.

### 3 Supplementary Figures & Tables

| Gene | Role | Function |
|---|---|---|
| Cbln1 | ligand | precerebellin; neuromodulatory functions |
| Cxcl14 | ligand | cytokine |
| Crhbp | receptor | Binds CRF |
| Gabra1 | receptor | gamma-aminobutyric acid (GABA) receptor |
| Cbln2 | ligand | paralog for Cbln1 |
| Gpr165 | receptor | G Protein-Coupled Receptor |
| Glra3 | receptor | Glycine receptors |
| Gabrg1 | receptor | GABAA receptors subunit |
| Adora2a | receptor | adenosine receptor of A2A subtype |
| Vgf | ligand | neuropeptide precursor |
| Scg2 | ligand | cytokine; neuroendocrine secretory protein |
| Cartpt | ligand | preproprotein for neuropeptides |
| Tac2 | ligand | tachykinin 2 |

Table S1: List of genes used as ligand and receptors in modeling and are not contained in the database Ramilowski *et al.* (2015). The roles and functions of the gene products are mostly obtained from `https://www.genecards.org/`.

| dataset | condition | selected values | outer CV test animal/replicate |
|---|---|---|---|
| MERFISH hypothalamus data | 84 | 0.0001 | 1,2,3,4 |
| STARmap mPFC data | 127 | 0.0001 | 1 |
| | | 0.00024 | 2 |
| | | 0.00147 | 3 |
| MERFISH U-2 OS cells | 38 | 0.02154 | B1 |
| | | 0.31623 | B2,B3 |
| | 74 | 0.02154 | B1 |
| | | 0.31623 | B2,B3 |
| | 181 | 0.31623 | B1,B2,B3 |
| | 364 | 0.31623 | B1,B2 |
| | | 0.05275 | B3 |
| | 693 | 0.02154 | B1 |
| | | 0.31623 | B2 |
| | | 0.05275 | B3 |

Table S2: Selected parameter values for the baseline model from the inner loop of the nested CV strategy for different datasets under different conditions (number of responses). The parameter for selection is alpha (penalty) of the lasso regression model.

| cell type | condition | selected values | outer CV test animal/replicate |
|---|---|---|---|
| Inhibitory | with neighborhood info | 10, soft | 1,2,3,4 |
| | no neighborhood info | 10, soft | 1,2,3,4 |
| Excitatory | with neighborhood info | 8, soft | 1,2 |
| | | 10, soft | 3,4 |
| | no neighborhood info | 10, soft | 1,2,3,4 |
| Astrocyte | with neighborhood info | 4, soft | 1,2 |
| | | 3, hard | 3,4 |
| | no neighborhood info | 3, soft | 1 |
| | | 4, hard | 2 |
| | | 3, hard | 3 |
| | | 5, hard | 4 |
| OD Mature 2 | with neighborhood info | 3, soft | 1,2 |
| | | 4, soft | 3 |
| | | 3, hard | 4 |
| | no neighborhood info | 3, hard | 1,2 |
| | | 4, hard | 3 |
| | | 5, hard | 4 |
| Endothelial 1 | with neighborhood info | 1, soft | 1,2 |
| | | 2, hard | 3,4 |
| | no neighborhood info | 2, hard | 1,2,3,4 |
| OD Immature 1 | with neighborhood info | 1, soft | 1,2 |
| | | 2, soft | 3,4 |
| | no neighborhood info | 2, soft | 1,2,3,4 |
| OD Mature 1 | with neighborhood info | 1, soft | 1,2,3,4 |
| | no neighborhood info | 1, soft | 1,2,3,4 |
| Microglia | with neighborhood info | 1, soft | 1,2,3,4 |
| | no neighborhood info | 1, soft | 1,2,3 |
| | | 2, soft | 4 |
| STARmap mPFC excitatory cells | with neighborhood info | 1, soft | 1,2,3 |
| | no neighborhood info | 1, soft | 1,2,3 |
| MERFISH U-2 OS cells* | with neighborhood info | 1, soft | B1,B2,B3 |
| | no neighborhood info | 1, soft | B1,B2,B3 |

Table S3: Selected parameter values for MESSI from the inner loop of the nested CV strategy for different cell types (datasets) under different conditions. Parameters for selection are 1) the number of experts 2) whether using “soft” or “hard” weighting among experts. See *Supplementary 1.1.4* for the difference between “soft” and “hard”. The first 8 cell types are from the MERFISH hypothalamus dataset. When the number of expert equals to 1, “soft” is the same as “hard” and thus only (1, soft) is listed. *: Models with different numbers of responses all result in the same parameter values.

| Cell type | Naive | Parenting | Virgin Parenting |
|---|---|---|---|
| Inhibitory | 9 | 8 | 8 |
| Excitatory | 8 | 5 | 5 |
| Astrocyte | 5 | 2 | 3 |
| OD Mature 2 | 5 | 2 | 2 |
| Endothelial 1 | 2 | 1 | 1 |
| OD Immature 1 | 2 | 1 | 1 |
| OD Mature 1 | 1 | 1 | 1 |
| Microglia | 1 | 1 | 1 |

Table S4: Number of experts used for naive and behavioral (parenting or virgin parenting) models for the comparison between naive and behavioral models on predicting behavioral samples of the MERFISH hypothalamus dataset.

| Cell type | Naive | Parenting | Virgin Parenting |
|---|---|---|---|
| Inhibitory | 9 | 8 | 8 |
| Excitatory | 8 | 8 | 8 |

Table S5: Number of experts used for naive and behavioral (parenting or virgin parenting) models trained on all available animals of the MERFISH hypothalamus dataset. These models are used for inference purposes.

| | Oxtr P | Oxtr VP |
|---|---|---|
| expert | | |
| 0.0 | 0.561724 | 0.512215 |
| 1.0 | 0.268403 | 0.324823 |
| 2.0 | 0.563461 | 0.381107 |
| 3.0 | 0.671636 | 0.693290 |
| 4.0 | 0.383936 | 0.378401 |
| 5.0 | 0.997566*** | 0.333738 |
| 6.0 | 0.548461 | 0.427870 |
| 7.0 | 0.301516 | 0.628811*** |

Table S6: Mean of the expression (raw count) of oxytocin receptor (Oxtr) in different experts (clusters of cells) in excitatory cells of the parenting (P) or virgin parenting (VP) animals of the MERFISH hypothalamus data. Expert 5 in parenting and expert 7 in virgin parenting, where both oxytocin and neighboring oxytocin show strong impact as shown in Figure 7, also have the largest and the second largest mean expression value of oxytocin receptor. ***: Oxtr in the expert is significantly higher than the other excitatory cells of the same behavior with a p-value lower than 1e-3 (one-sided Mann-Whitney U-test).

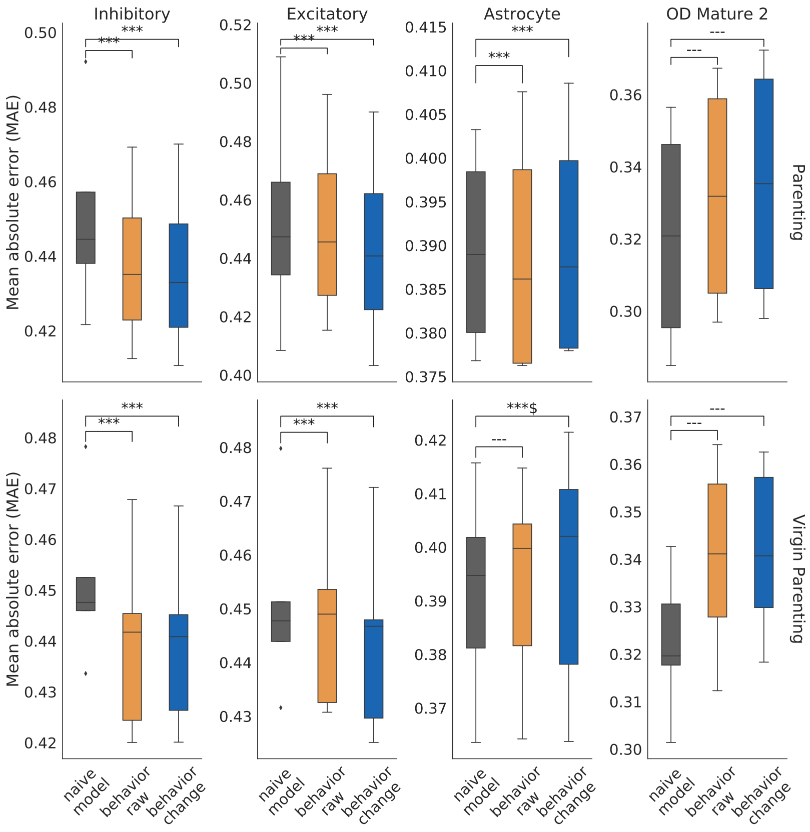

Figure S1: Comparison of naive and behavior models. Top: when the behavior is parenting. Bottom: when the behavior is virgin parenting. Results are presented for the 4 major cell types. Naive model - predictions based on the naive model. Behavior raw - predictions based on learning from the raw values in the corresponding behavioral samples. Behavior change - predictions based on learning from the raw values in the corresponding behavioral samples subtracted by the predictions from the naive model. See *Supplementary Methods* for details. ***: p-value below 1e-3; **: p-value below 1e-2; *: p-value below 5e-2; ---: p-value larger than 5e-2; $: more than half of the CV groups show non-significant improvement when using behavioral models.

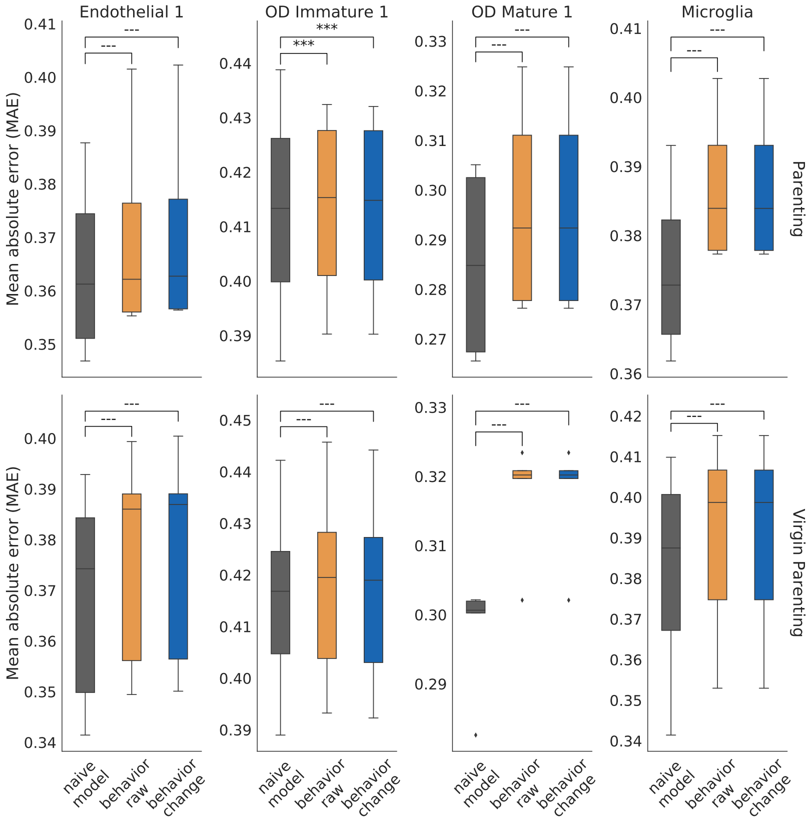

Figure S2: Comparison of naive and behavior models. Top: when the behavior is parenting. Bottom: when the behavior is virgin parenting. Results are presented for the 4 smaller cell types. Naive model - predictions based on the naive model. Behavior raw - predictions based on learning from the raw values in the corresponding behavioral samples. Behavior change - predictions based on learning from the raw values in the corresponding behavioral samples subtracted by the predictions from the naive model. See *Supplementary Methods* for details. ***: p-value below 1e-3; **: p-value below 1e-2; *: p-value below 5e-2; ---: p-value larger than 5e-2; $: more than half of the CV groups show non-significant improvement when using behavioral models.

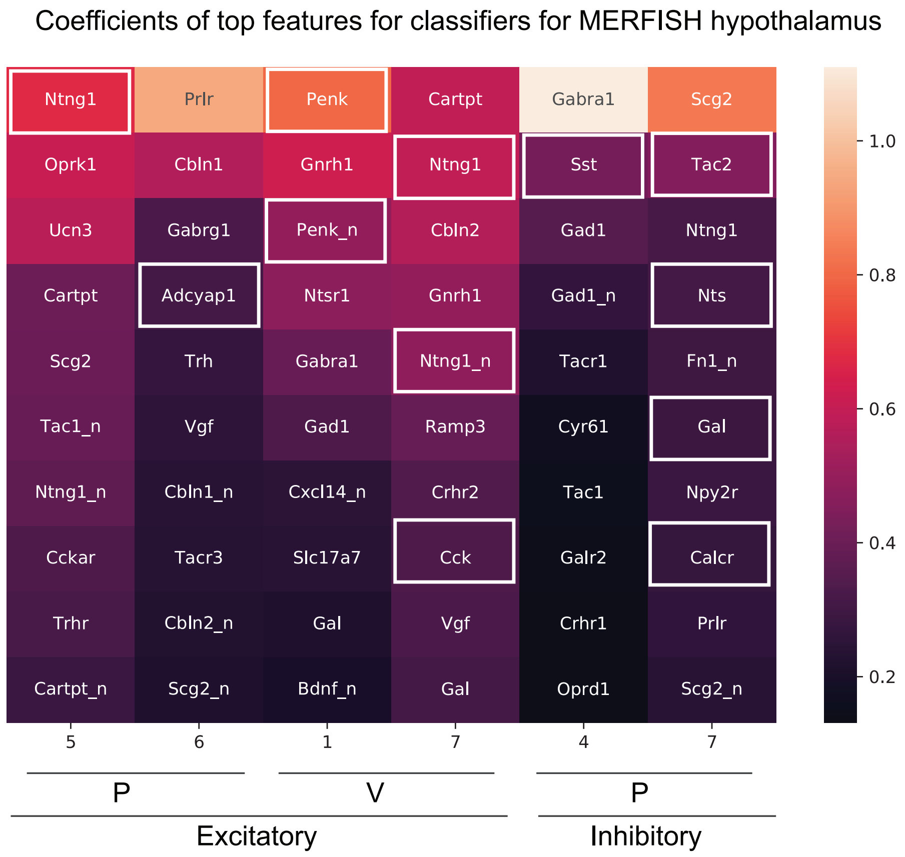

Figure S3: Top features (gene names; the ones with suffix "_n" are expressed by the neighbors) ranked by the coefficient (represented by different colors), for classifiers trained for the MERFISH hypothalamus data for two cell types in different behavioral samples (P: parenting females; V: virgin parenting females). Each column represents the top 10 features for an expert. The ones with white blocks are also found as differential expressed genes of corresponding clusters in Moffitt *et al.* (2018).

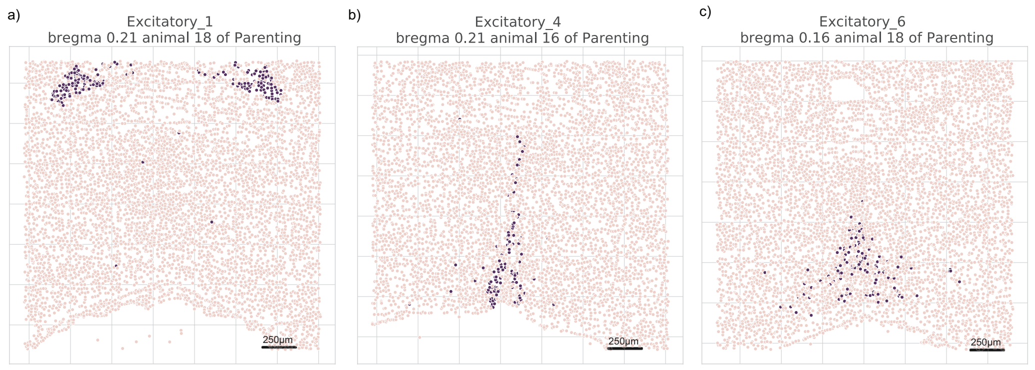

Figure S4: Spatial location of the cells on an example bregma for excitatory cells in parenting samples. a): location of cells assigned to expert 1; b): location of cells assigned to expert 4; c): location of cells assigned to expert 6

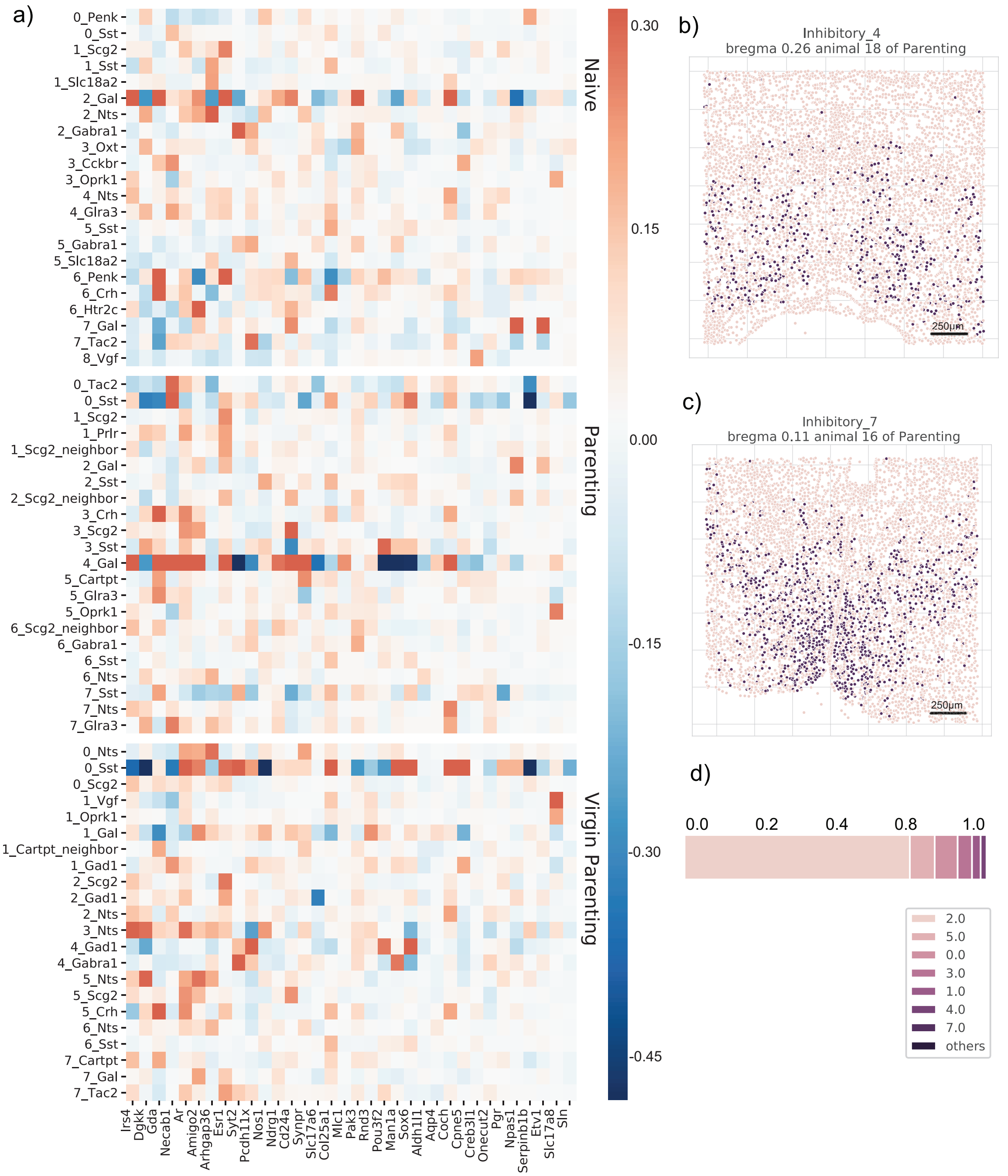

Figure S5: MESSI reveals changes in key signaling molecules and relevant signaling networks activated upon experience. Left a): Coefficients for top signaling molecules in a subset of the MESSI experts of inhibitory cells for different behaviors (Y axis) for several response genes (X axis). See *Supporting Methods* for selection of top features. Right: Cells assigned to specific MESSI experts. b), c): spatial location of the cells on an example bregma; d): the proportion of cells from expert 4 in parenting assigned to different experts by the naive model; "_neighbor": feature expressed by the neighboring cells; The feature names are prefixed by the expert number in a)

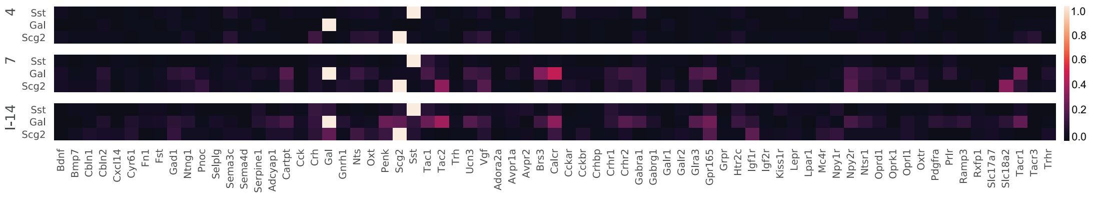

Figure S6: Pairwise correlations between Sst, Gal, Scg2, and other signaling genes (expressed by cells themselves). We calculate the correlations for three subgroups of cells, 4: inhibitory expert 4 of parenting; 7: inhibitory expert 7 of parenting; I-14: inhibitory cluster assigned by Moffitt *et al.* (2018). Note that expert 7 overlaps with I-14. Gal correlates strongly with other features such as bombesin receptor subtype 3 (Brs3) and calcitonin receptor (Calcr) in both expert 7 and the cluster I-14. In comparison, Sst does not have obvious collinearity with any other ligand or receptors.

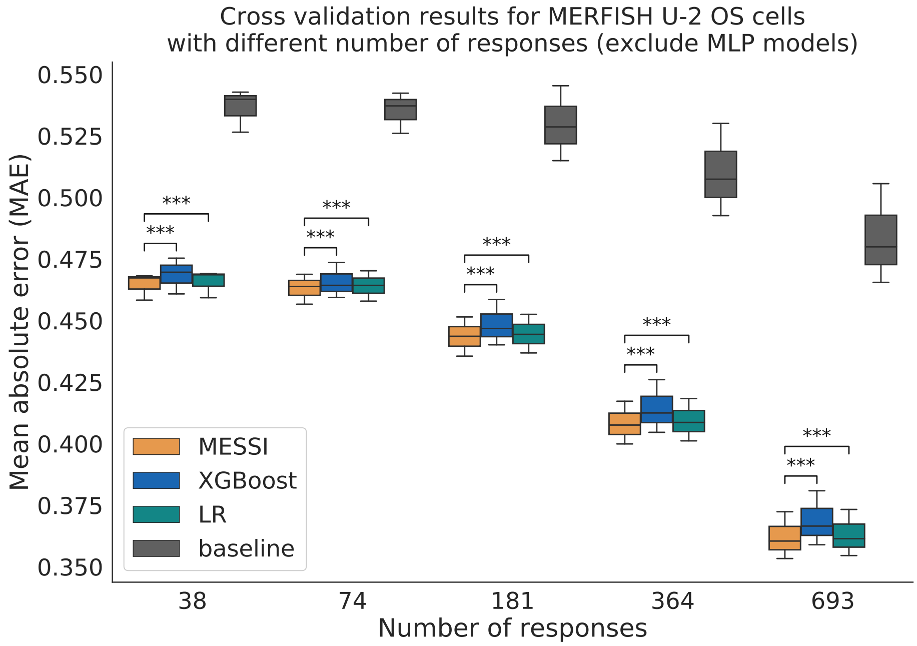

Figure S7: CV results for the MERFISH U-2 OS data from top performing models and the baseline using different numbers of response genes. p-values indicate the significance level of MESSI outperforming the comparison model. Based on Table S3, here MESSI was selected from the inner loops to have only 1 single expert and thus is equivalent to MROTS (thus not included in this plot). MLP models' performance are very different from the top tiers and thus excluded from this plot. See Figure S10 for an example. LR: Linear regression; MLP single: multi-layer perceptron (MLP) with single output node; MLP multi: multi-layer perceptron (MLP) with multiple output nodes; MROTS: Multiple-output Regression with Output and Task Structures. ***: p-value below 1e-3; **: p-value below 1e-2; *: p-value below 5e-2

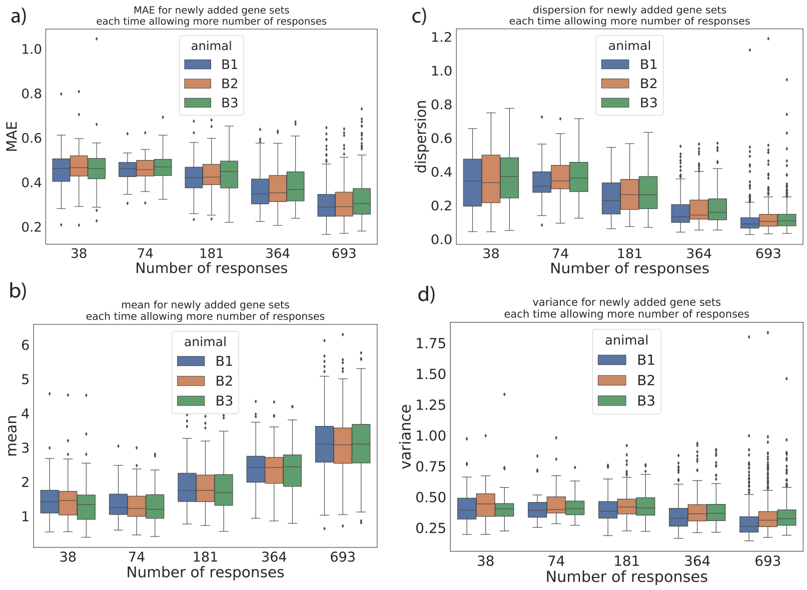

Figure S8: MAE (averaged over cells only) and statistics of new genes added each time when number of responses increases (for example, genes shown for the model of 74 responses are those not used for the 38 responses model and genes for the model of 181 responses are those not used for the 74 responses model etc.) of the MERFISH U-2 OS dataset. Dispersion is defined as variance/mean.

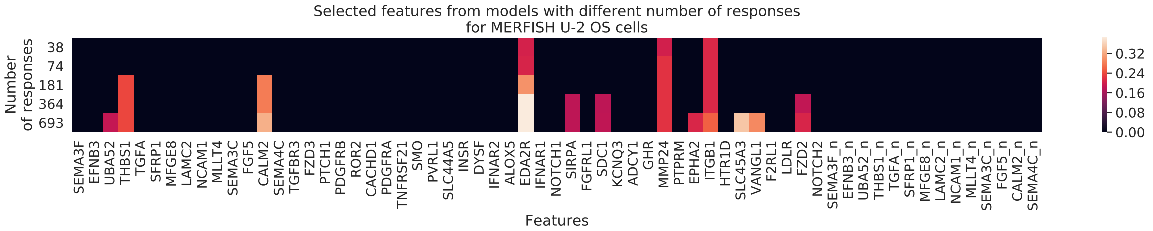

Figure S9: Top features selected for models using different numbers of response genes for the MERFISH U-2 OS dataset. See *Supplementary 1.4.4* for the selection criteria. _n: ligand expressed by the neighbors

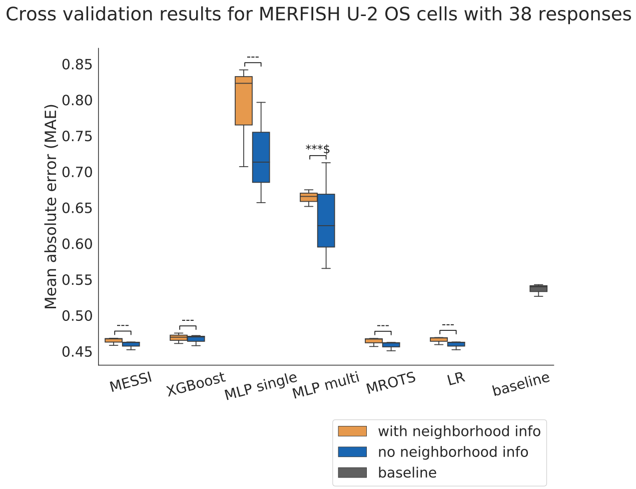

Figure S10: Comparison between the CV results when using neighborhood information and not using neighborhood information for different models on the MERFISH U-2 OS cells when using 38 response variables. Based on Table S3, here MESSI was selected from the inner loops to have only 1 single expert and thus is equivalent to MROTS. LR: Linear regression; MLP single: multi-layer perceptron (MLP) with single output node; MLP multi: multi-layer perceptron (MLP) with multiple output nodes; MROTS: Multiple-output Regression with Output and Task Structures. ***: p-value below 1e-3; **: p-value below 1e-2; *: p-value below 5e-2; ---: p-value larger than 5e-2; $: more than half of the CV groups show non-significant improvement when using neighborhood information.

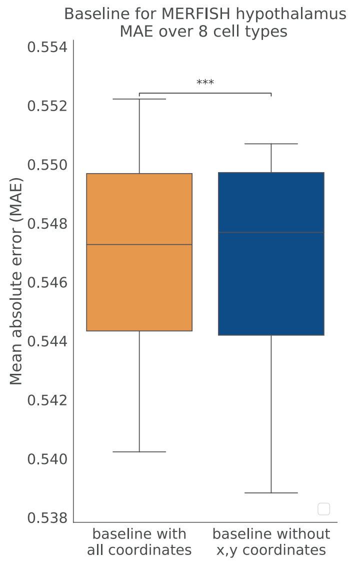

Figure S11: CV results comparing baseline models using or not using x, y coordinates as features for the MERFISH hypothalamus dataset. The p-value indicates the significance level of the model using x, y coordinates outperforming the one not using. ***: p-value below 1e-3.

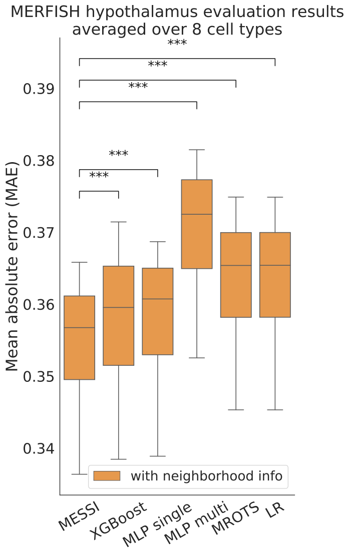

Figure S12: Comparison between the CV results of MESSI and other models averaged over all cells of 8 cell types when using neighborhood information of the MERFISH hypothalamus data. p-values indicate the significance levels of MESSI outperforming the comparison model. LR: Linear regression; MLP single: multi-layer perceptron (MLP) with single output node; MLP multi: multi-layer perceptron (MLP) with multiple output nodes; MROTS: Multiple-output Regression with Output and Task Structures. ***: p-value below 1e-3

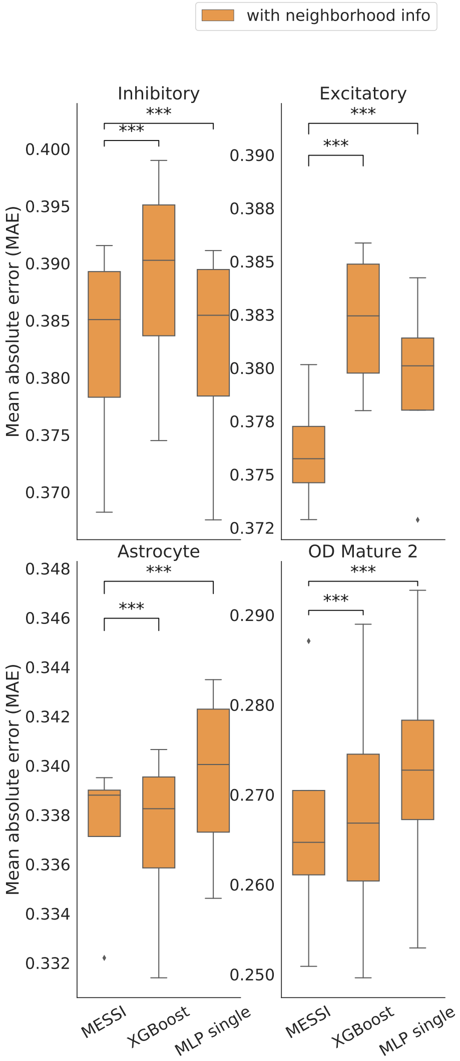

Figure S13: Comparison between the CV results of MESSI and other models when using neighborhood information for the major cell types of the MERFISH hypothalamus data. p-values indicate the significance levels of MESSI outperforming the comparison model. LR: Linear regression; MLP single: multi-layer perceptron (MLP) with single output node; MLP multi: multi-layer perceptron (MLP) with multiple output nodes; MROTS: Multiple-output Regression with Output and Task Structures. ***: p-value below 1e-3

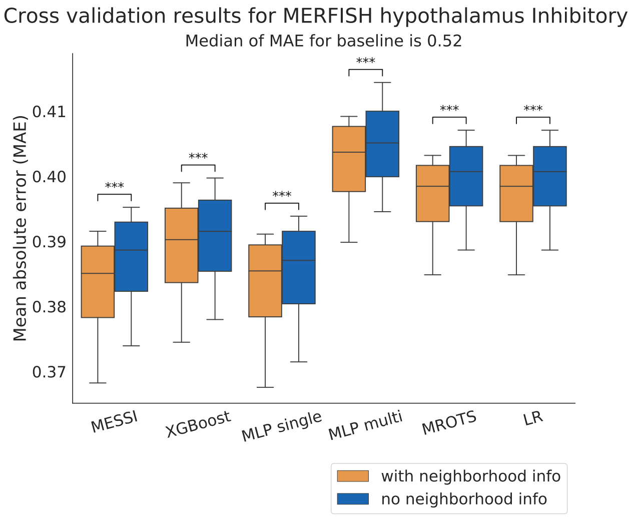

Figure S14: Comparison between the CV results when using neighborhood information and not using neighborhood information for different models on the Inhibitory cell type of the MERFISH hypothalamus data. LR: Linear regression; MLP single: multi-layer perceptron (MLP) with single output node; MLP multi: multi-layer perceptron (MLP) with multiple output nodes; MROTS: Multiple-output Regression with Output and Task Structures. ***: p-value below 1e-3; **: p-value below 1e-2; *: p-value below 5e-2; ---: p-value larger than 5e-2; $: more than half of the CV groups show non-significant improvement when using neighborhood information.

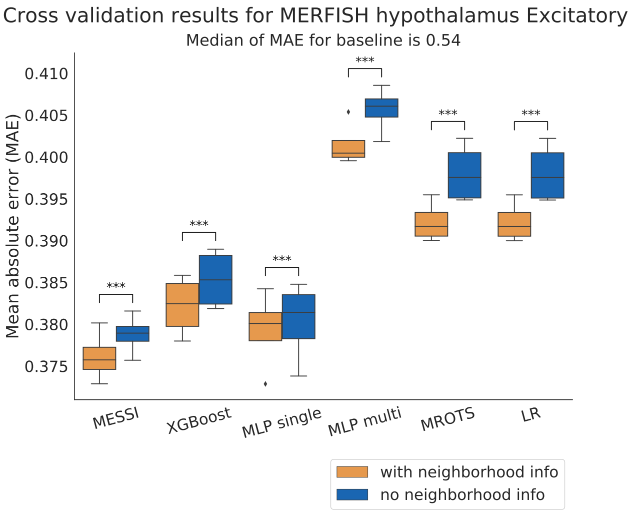

Figure S15: Comparison between the CV results when using neighborhood information and not using neighborhood information for different models on the Excitatory cell type of the MERFISH hypothalamus data. LR: Linear regression; MLP single: multi-layer perceptron (MLP) with single output node; MLP multi: multi-layer perceptron (MLP) with multiple output nodes; MROTS: Multiple-output Regression with Output and Task Structures. ***: p-value below 1e-3; **: p-value below 1e-2; *: p-value below 5e-2; ---: p-value larger than 5e-2; $: more than half of the CV groups show non-significant improvement when using neighborhood information.

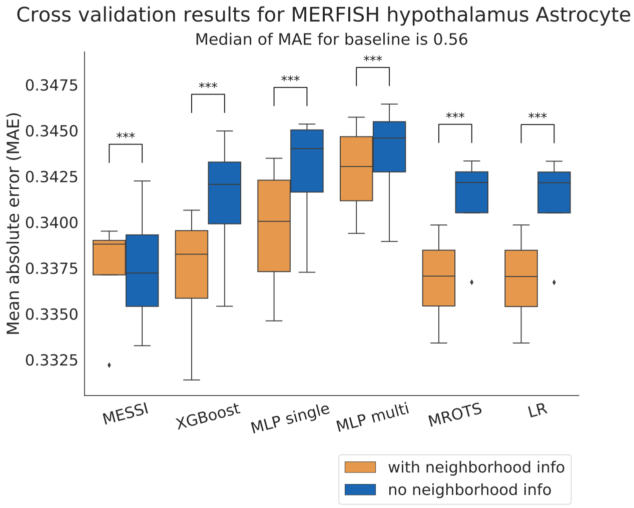

Figure S16: Comparison between the CV results when using neighborhood information and not using neighborhood information for different models on the Astrocyte cell type of the MERFISH hypothalamus data. LR: Linear regression; MLP single: multi-layer perceptron (MLP) with single output node; MLP multi: multi-layer perceptron (MLP) with multiple output nodes; MROTS: Multiple-output Regression with Output and Task Structures. ***: p-value below 1e-3; **: p-value below 1e-2; *: p-value below 5e-2; ---: p-value larger than 5e-2; $: more than half of the CV groups show non-significant improvement when using neighborhood information.

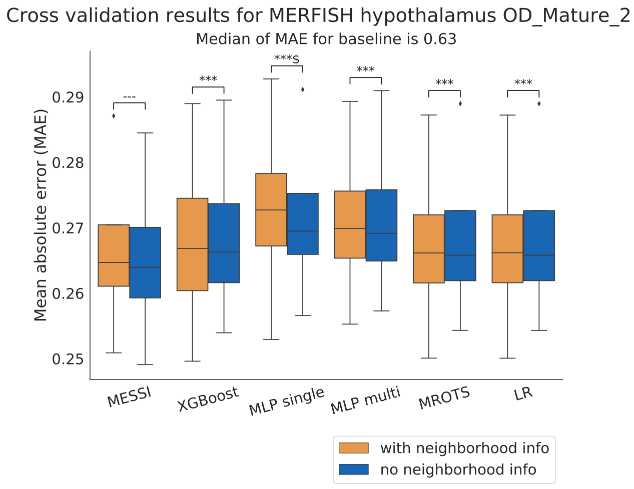

Figure S17: Comparison between the CV results when using neighborhood information and not using neighborhood information for different models on the OD Mature 2 cell type of the MERFISH hypothalamus data. LR: Linear regression; MLP single: multi-layer perceptron (MLP) with single output node; MLP multi: multi-layer perceptron (MLP) with multiple output nodes; MROTS: Multiple-output Regression with Output and Task Structures. ***: p-value below 1e-3; **: p-value below 1e-2; *: p-value below 5e-2; ---: p-value larger than 5e-2; $: more than half of the CV groups show non-significant improvement when using neighborhood information.

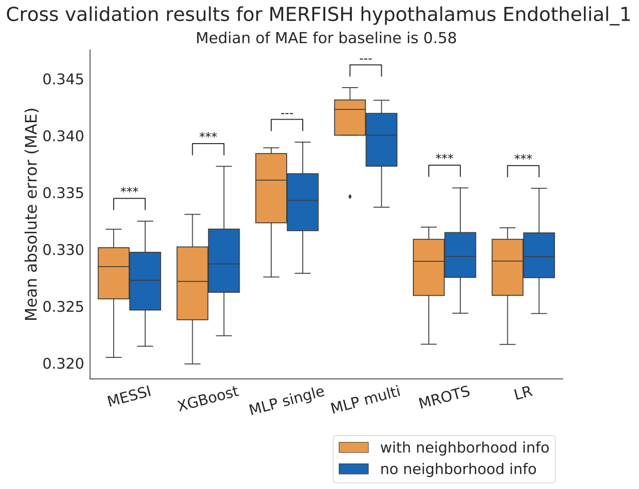

Figure S18: Comparison between the CV results when using neighborhood information and not using neighborhood information for different models on the Endothelial 1 cell type of the MERFISH hypothalamus data. LR: Linear regression; MLP single: multi-layer perceptron (MLP) with single output node; MLP multi: multi-layer perceptron (MLP) with multiple output nodes; MROTS: Multiple-output Regression with Output and Task Structures. ***: p-value below 1e-3; **: p-value below 1e-2; *: p-value below 5e-2; ---: p-value larger than 5e-2; $: more than half of the CV groups show non-significant improvement when using neighborhood information.

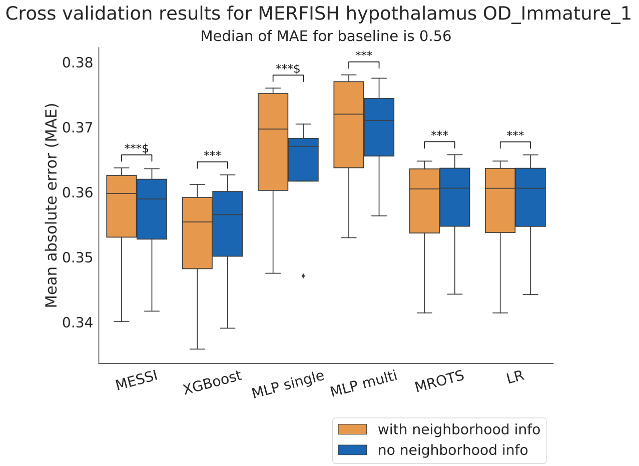

Figure S19: Comparison between the CV results when using neighborhood information and not using neighborhood information for different models on the OD Immature 1 cell type of the MERFISH hypothalamus data. LR: Linear regression; MLP single: multi-layer perceptron (MLP) with single output node; MLP multi: multi-layer perceptron (MLP) with multiple output nodes; MROTS: Multiple-output Regression with Output and Task Structures. ***: p-value below 1e-3; **: p-value below 1e-2; *: p-value below 5e-2; ---: p-value larger than 5e-2; $: more than half of the CV groups show non-significant improvement when using neighborhood information.

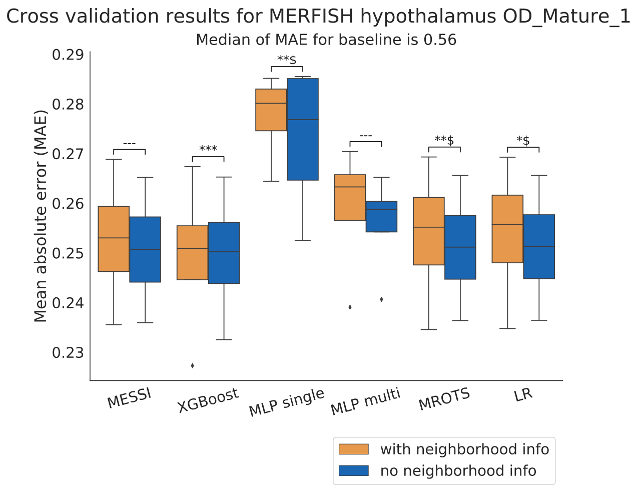

Figure S20: Comparison between the CV results when using neighborhood information and not using neighborhood information for different models on the OD Mature 1 cell type of the MERFISH hypothalamus data. LR: Linear regression; MLP single: multi-layer perceptron (MLP) with single output node; MLP multi: multi-layer perceptron (MLP) with multiple output nodes; MROTS: Multiple-output Regression with Output and Task Structures. ***: p-value below 1e-3; **: p-value below 1e-2; *: p-value below 5e-2; ---: p-value larger than 5e-2; $: more than half of the CV groups show non-significant improvement when using neighborhood information.

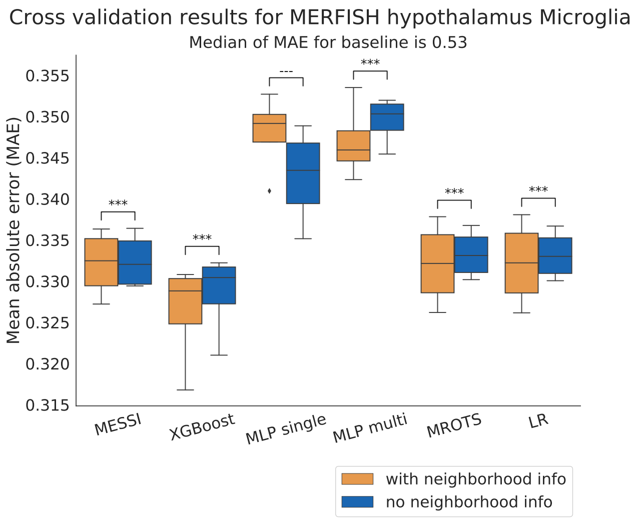

Figure S21: Comparison between the CV results when using neighborhood information and not using neighborhood information for different models on the Microglia cell type of the MERFISH hypothalamus data. LR: Linear regression; MLP single: multi-layer perceptron (MLP) with single output node; MLP multi: multi-layer perceptron (MLP) with multiple output nodes; MROTS: Multiple-output Regression with Output and Task Structures. ***: p-value below 1e-3; **: p-value below 1e-2; *: p-value below 5e-2; ---: p-value larger than 5e-2; $: more than half of the CV groups show non-significant improvement when using neighborhood information.

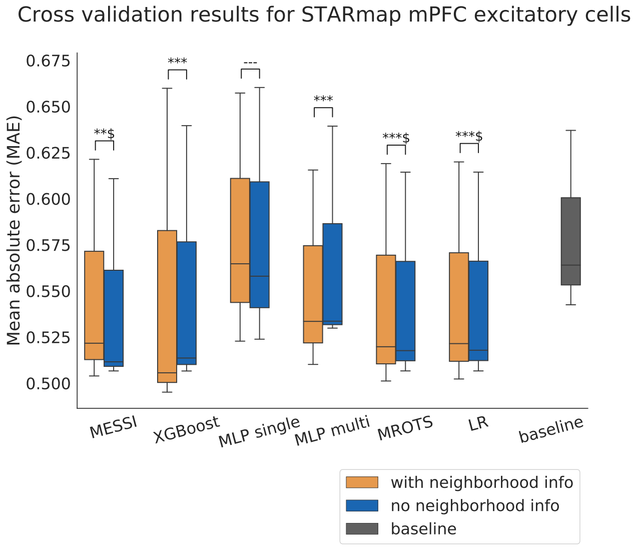

Figure S22: Comparison between the CV results when using neighborhood information and not using neighborhood information for different models on the excitatory cell type of the STARmap data. Based on Table S3, here MESSI was selected from the inner loops to have only 1 single expert and thus is equivalent to MROTS. LR: Linear regression; MLP single: multi-layer perceptron (MLP) with single output node; MLP multi: multi-layer perceptron (MLP) with multiple output nodes; MROTS: Multiple-output Regression with Output and Task Structures. ***: p-value below 1e-3; **: p-value below 1e-2; *: p-value below 5e-2; ---: p-value larger than 5e-2; $: more than half of the CV groups show non-significant improvement when using neighborhood information.

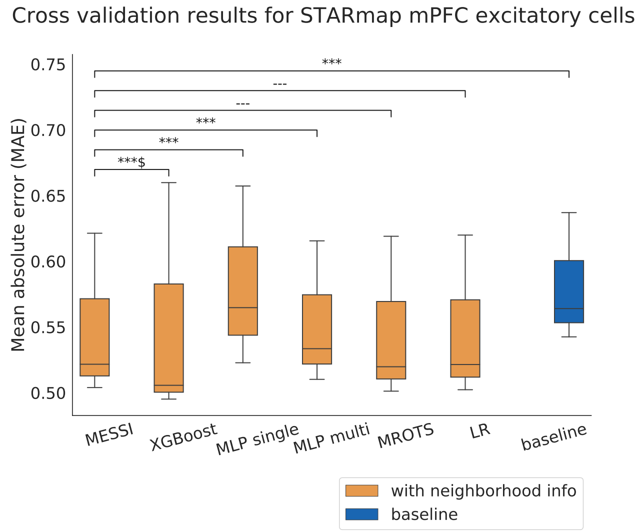

Figure S23: Comparison between the CV results of MESSI and other models on the excitatory cell type of the STARmap data when using neighborhood information. p-values indicate the significance levels of MESSI outperforming the comparison model. Based on Table S3, here MESSI was selected from the inner loops to have only 1 single expert and thus is equivalent to MROTS. LR: Linear regression; MLP single: multi-layer perceptron (MLP) with single output node; MLP multi: multi-layer perceptron (MLP) with multiple output nodes; MROTS: Multiple-output Regression with Output and Task Structures. ***: p-value below 1e-3; **: p-value below 1e-2; *: p-value below 5e-2; ---: p-value larger than 5e-2; $: more than half of the CV groups show non-significant improvement when using MESSI.
